# Comprehensive detection and characterization of human druggable pockets through novel binding site descriptors

**DOI:** 10.1101/2024.03.14.584971

**Authors:** Arnau Comajuncosa-Creus, Guillem Jorba, Xavier Barril, Patrick Aloy

## Abstract

Druggable pockets are protein regions that have the ability to bind organic small molecules, and their characterization is essential in target-based drug discovery. However, strategies to derive pocket descriptors are scarce and usually exhibit limited applicability. Here, we present PocketVec, a novel approach to generate pocket descriptors for any protein binding site of interest through the inverse virtual screening of lead-like molecules. We assess the performance of our descriptors in a variety of scenarios, showing that it is on par with the best available methodologies, while overcoming some important limitations. In parallel, we systematically search for druggable pockets in the folded human proteome, using experimentally determined protein structures and AlphaFold2 models, identifying over 32,000 binding sites in more than 20,000 protein domains. Finally, we derive PocketVec descriptors for each small molecule binding site and run an all-against-all similarity search, exploring over 1.2 billion pairwise comparisons. We show how PocketVec descriptors facilitate the identification of druggable pocket similarities not revealed by structure- or sequence-based comparisons. Indeed, our analyses unveil dense clusters of similar pockets in distinct proteins for which no inhibitor has yet been crystalized, opening the door to strategies to prioritize the development of chemical probes to cover the druggable space.

## Introduction

Ligand binding sites are protein regions that interact with other biochemical entities such as peptides or organic small molecules. The binding process eventually results in a selective modulation of the protein function. Indeed, one of the most successful strategies in conventional drug discovery is to identify, based on the high-resolution three-dimensional structure of binding sites, small molecules that activate or inhibit a protein associated with a disease^1^.

Alongside the increasing number of available protein structures in the Protein Data Bank (PDB)^2^, structure-based approaches have become a crucial computational framework in early stages of drug development^3, 4^. By focusing on ligand binding sites, such strategies enable a rational design and optimization of drugs and reduce the probability of failure of those compounds that reach clinical trials^5^. Protein-small molecule docking is among the most popular structure-based strategies to predict drug-target interactions, and it aims at finding the optimal location and conformation of a given ligand with respect to the receptor binding site^6^. Molecular docking has been successfully applied in proteome-scale studies (e.g. reverse screening^7–9^), but the target-dependent nature of scoring functions prevents the direct comparisons of docking results across different proteins and protein families. Indeed, the design of a universal docking scoring function still remains a challenge^10, 11^. Consequently, alternative strategies such as reverse pharmacophore screening, binding site similarity assessment or interaction fingerprint comparison are often employed in proteome-wide analyses^12^.

Most of these approaches require the detailed characterization of protein binding sites in a machine-readable format suitable for computational applications, which reasonably allows for the possibility of borrowing featurization techniques from related fields. In fact, characterizing small molecules through numerical vectors encoding topological or physicochemical properties is a very common strategy in cheminformatics, and sets the stage for many drug discovery projects founded on the small-molecule similarity principle^13–15^. Likewise, descriptors for larger molecules, such as protein targets, can also be derived, usually gathering features from their amino acid sequences^16^. However, the exploitation of structural data offers a complementary perspective to create protein descriptors and is therefore more promising than the treatment of protein sequences alone. Indeed, biophysical interactions between proteins and ligands occur in very specific areas of protein surfaces (i.e. binding sites) and involve a limited set of residues, which has driven the development of structure-based protein descriptors focused on these particular regions^17^.

Pocket descriptors are commonly classified according to the underlying binding site representation they consider, often based on binding site residues (e.g. FuzCav^18^, SiteAlign^19^), pocket surfaces (e.g. MaSIF^20^) or explicit interactions with bound ligands or probes (e.g. KRIPO^21^, TIFP^22^, BioGPS^23^). In addition, and together with the rising interest in deep learning applications in drug discovery^24^, novel data-driven approaches have been designed to derive pocket descriptors borrowing techniques from computer vision (e.g. DeeplyTough^25^, BindSiteS-CNN^26^).

Apart from the inherent characterization of binding sites, pocket descriptors provide an excellent means to estimate binding site similarity, which is thus simplified into straightforward vector distance measurements. Binding site comparisons (*aka* pocket matching) have emerged as a promising methodology to move away from the ‘one drug-one target-one disease’ paradigm^27^ by assessing complex studies involving multiple-target drug binding events^28–30^. Binding site similarity is reported to play an important role in the evaluation of ligand promiscuity^31^ and in the prediction of protein function, enabling the identification of similar binding sites in proteins having no sequence nor fold similarity^32^. Indeed, the detection of similar binding sites was helpful in several drug repurposing and polypharmacology studies^33–37^ and in the prediction of possible distant drug off-targets^38^. In addition, encoding pockets as numerical descriptors entails the possibility of integrating them in a unified framework together with a rich portrait of biochemical entities described in a common vectorial format, such as small molecules, cell lines or diseases^13^. For instance, chemogenomic studies are often addressed by the combination of protein descriptors and molecular fingerprints, usually referred to as proteochemometric (PCM) approaches^39, 40^.

However, existing methods to generate pocket descriptors exhibit several intrinsic limitations. One of their main drawbacks is the need of co-crystallized ligands to effectively recognize the most relevant biophysical interactions occurring in the binding site, which restricts the applicability domain of such methods to *holo* structures^21, 22^. Another important issue related with pocket descriptors is the handcrafted nature of considered binding site representations, often selecting parameters based on specific datasets and performing poorly when used in more general and diverse scenarios^41^. Moreover, several strategies also rely on alignment-dependent comparisons, which makes them particularly useful to provide significant insights into the underlying patterns rationalizing binding site similarity, but also come together with an increased computational cost^19^. In addition, those approaches built upon deep learning algorithms also suffer from lack of interpretability, a well-known problem in the field^42–44^. Finally, the availability of three-dimensional protein structures has traditionally been the main limiting factor in structure-based drug discovery, but this is no longer the case. In the era of accurate protein structure prediction^45–47^, where exhaustive collections of predicted structures are available for both relevant organisms^48^ and sequences derived from metagenomic studies^49^, the structural characterization of proteins is now feasible for essentially any protein sequence of interest. Accordingly, the use of pocket descriptors opens the possibility of characterizing complete proteomes and charting the pocket space in a similar way molecular fingerprints enable the exploration of the chemical space of small molecules^50–52^.

To partially overcome the aforementioned limitations of existing pocket descriptors, we exploit the assumption that similar pockets bind similar ligands, which should result in similar rankings in a structure-based virtual screening of small molecules. Indeed, Govindaraj and Brylinski^53^ showed that docking scores tended to be more correlated in pockets binding to chemically similar ligands than in pockets binding to dissimilar ligands. This opens the possibility of estimating binding site similarity on the basis of docking rankings and enrichments, as explored by Schmidt and co-workers in their analysis of the human kinome^54^. Moreover, inverse virtual screening (i.e. the screening of a set of targets for a query ligand) has been recently applied to distinguish nucleotide and heme-binding sites from a control set of pockets^55^. In view of these results, we hypothesized that virtual screening could represent a promising strategy to generate pocket descriptors.

Here, we present PocketVec, a novel strategy to generate interpretable and fixed-length protein binding site descriptors based on the assumption that similar pockets bind similar ligands. Our approach is built upon inverse virtual screening, i.e. the prioritization of a given set of small molecules is expected to be more correlated between similar pockets than between dissimilar ones. We implement and assess the accuracy of our method and the derived pocket descriptors on several predefined benchmark sets. Additionally, we use bound ligands and pocket detection algorithms to comprehensively identify drug binding pockets in experimentally determined and AF2 predicted structures in the human proteome, and derive PocketVec descriptors for all identified pockets. We finally use PocketVec descriptors to exhaustively compare all pockets found in experimental and AF2 structures, to show the complementarity with other sequence and structure-based approaches and to demonstrate its potential to find and characterize similar binding sites in unrelated proteins.

## Methodological development and implementation

It is known that similar proteins tend to bind similar ligands^56^, a principle behind many drug discovery projects^12, 57, 58^. We re-assessed the validity of this principle and found that, indeed, proteins from the same family (e.g. GPCRs) tend to have more similar active compounds than proteins from different families (**Fig S1**). However, globally dissimilar proteins showing similar physicochemical and shape properties in their druggable pockets may still bind with similar ligands, which reasonably translates into a more precise and general form of the chemogenomics principle: similar pockets bind similar ligands^59^. PocketVec builds on this observation to generate novel vector-type descriptors for characterizing protein small molecule binding pockets.

Instead of directly characterizing the shape and physicochemical environment of the protein cavities, we rely on a predefined set of small molecules and assess their potential binding to a given pocket. More specifically, given a three-dimensional protein structure, we first identify possible druggable pockets and we then use computational docking strategies to assess the potential binding of the small molecules. The resulting docking scores are then translated into rankings, which are finally stored in a vector-type format. In this way, each bit of the vector represents the ranking of a predefined molecule, illustrating how good it binds with the pocket of interest compared to all other molecules (**Fig 1a**). While the idea is conceptually pretty straightforward, its implementation requires a thorough assessment of the set of used molecules, the docking methodology and the benchmark strategy. The following sections describe our effort to evaluate and optimize each step of the procedure.

**Fig 1:**
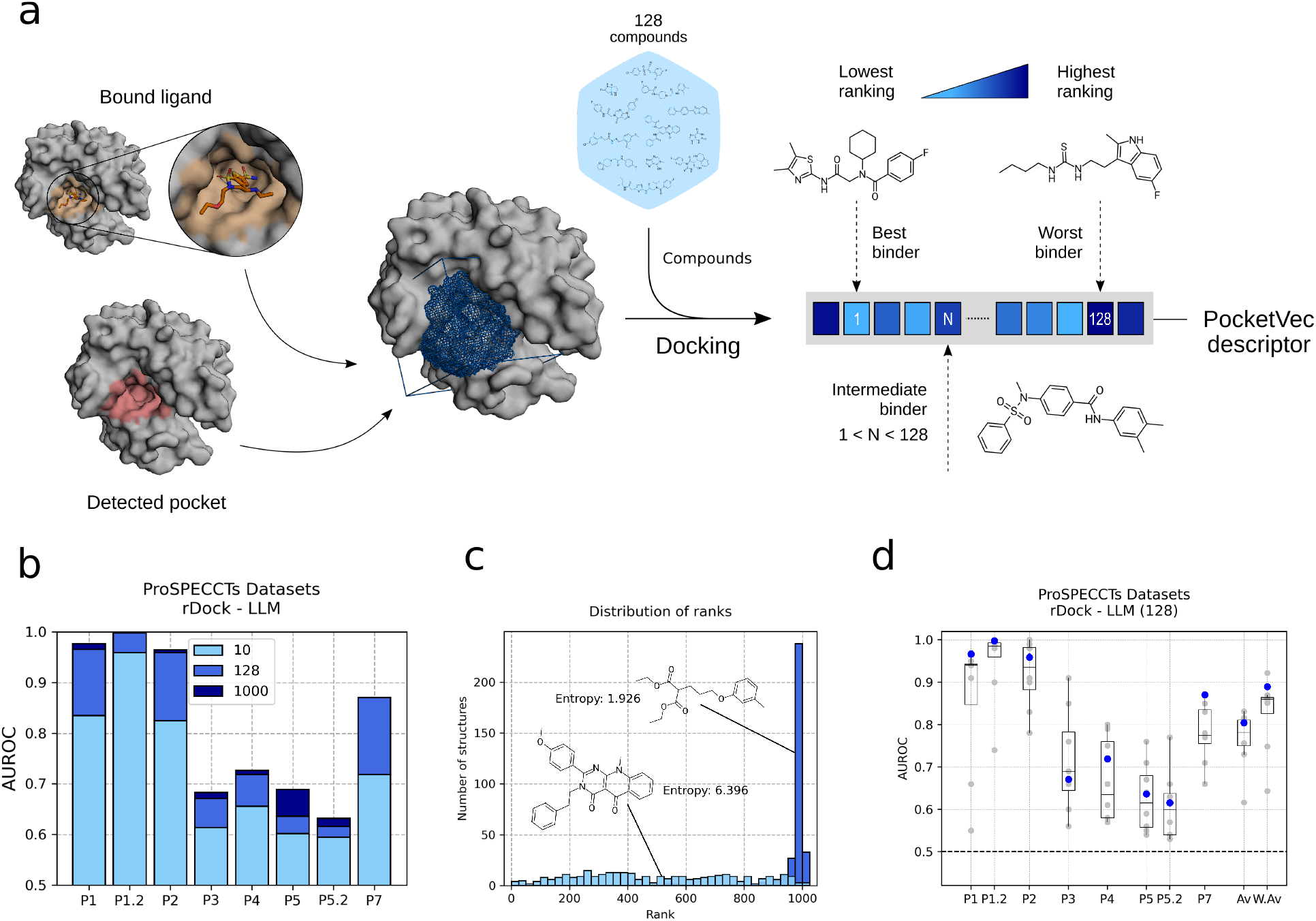
PocketVec methodological pipeline and benchmark results. **a)** Given a 3D protein structure, binding site locations are established by the presence of bound ligands or by means of pocket detection algorithms. A predefined set of compounds (128 lead-like molecules in the standard PocketVec pipeline) is docked against the pocket of interest. The corresponding docking scores are then converted into rankings and stored in a vector-type format that serves to characterize the pocket. We refer to those vectors as PocketVec descriptors. **b)** Bars indicating the performance (AUROC, y-axis) of our descriptors generated with a varying number of predefined molecules among ProSPECCTs datasets (x-axis, P6 and P6.2 not included. For further details, please see *Online Methods*). Bar color indicates the number of predefined compounds (10, 128 and the complete set of 1,000 lead-like molecules, sorted by entropy). These results correspond to the rigid docking (rDock) and LLM combination. All the other combinations with all possible numbers of predefined compounds (from 1 to complete sets) are shown in **Fig S7**. **c)** Predefined molecules with high and low entropy. The histograms depict the distribution (y-axis) of rankings (x-axis, bin width: 25) for the highest (sky blue) and lowest (dark blue) entropy lead-like molecules in ProSPECCTs P1. Their chemical structures are shown together with the corresponding entropy values. **d)** Performances (AUROC, y-axis) of distinct pocket descriptors among ProSPECCTs datasets (x-axis, P6 and P6.2 not included). Gray dots represent the individual performances of existing strategies to derive pocket descriptors (see **Table S1**). Box plots indicate median (middle line), 25th, 75th percentile (box), and max and min value within the 1.5*25th and 1.5*75th percentile range (whiskers). Blue dots indicate the performance of PocketVec descriptors (128 LLM and rDock rigid docking). Av. values represent the average performance among ProSPECCTs datasets for each individual method and W. Av. values weight the average value according to the number of pairs within each dataset.

### Selection of the methodological pipeline

To find the optimal set of compounds to develop our binding pocket descriptors, we tested two different types of molecules. On the one hand, we used the Glide chemically diverse collection of fragments^60, 61^, containing 667 compounds with molecular weights in the 50-200 g·mol^-1^ range (**Fig S2**). Additionally, we also selected 1,000 lead-like molecules (LLM) from the MOE v2019.01 dataset (Chemical Computing Group, Montreal, Canada), exhibiting molecular weights in the 200-450 g·mol^-1^ range (**Fig S2**).

We also assessed the performance of two well-established small molecule docking strategies. More specifically, we used rDock^62^ and SMINA^63^ to run rigid and flexible docking calculations, respectively, under the default parameters.

Finally, to determine the best combination of small molecules and docking methods we relied on ProSPECCTs, a collection of datasets aimed at evaluating the performance of pocket comparison approaches (including pocket descriptors) in a wide range of distinct scenarios^41^. In brief, ProSPECCTs comprises 10 datasets consisting of protein-ligand binding site pairs classified as either similar or dissimilar according to various criteria, including pairs of different structures of the same proteins, proteins harboring artificial mutations in their binding pockets or pairs of unrelated proteins that are able to bind chemically similar ligands (**Fig S3**).

Please, see the *Online Methods* section for a more detailed description of the methodological pipeline.

### PocketVec parameter selection and benchmark

For each ProSPECCTs dataset, we generated PocketVec descriptors for all ligand-defined protein binding sites, and evaluated pocket similarity on the basis of pairwise cosine distances between PocketVec descriptors: the lower the PocketVec distance, the higher the pocket similarity.

First, we obtained results for all possible combinations of docking strategies (rDock - rigid docking and SMINA - flexible docking) and compound collections (1,000 lead-like molecules and 667 fragments). We observed that, in general, the use of LLM and rDock rigid docking provided better results than other combinations (**Fig S4**). Positive docking scores (i.e. molecules that could not be accommodated in the binding pocket, see *Online Methods* for further details) leading to outlier rankings were rare with fragments but more frequent when using LLM (∼35% of structures had at least one outlier molecule in the rDock - LLM (1,000) combination, **Fig S5**). Thus, LLM showed a superior discriminative power with respect to the size of the pocket and, overall, provided higher ranking diversity among all ProSPECCTs pockets (**Fig S6**). In fact, we observed how LLM occupied a larger fraction of the binding sites than fragments, while rigid docking might have removed the noise created by similar rankings of the same ligand in different conformations, overall conferring a superior discrimination capacity to the rDock - LLM combination.

Our benchmarks also revealed that some molecules were systematically ranked as very weak binders and were thus not informative for the assessment of pocket similarity. Indeed, we realized that the use of ∼100-200 molecules was often sufficient to get competitive performances in most datasets. In order to optimize the set of predefined molecules to derive pocket descriptors, we selected those LLM and fragments presenting high ranking diversity across all ProSPECCTs datasets (see *Online Methods*). In brief, we calculated the Shannon’s entropy for each molecule within each dataset and, after intra-dataset normalization, we assigned each molecule an averaged entropy value, which enabled us to prioritize those molecules presenting high ranking diversity (high entropy). Indeed, performances obtained in ProSPECCTs datasets by the 128 most diverse molecules were fairly similar to the original ones using complete sets of compounds (**Fig 1b**, **Fig S7**). Reducing the number of docked molecules (from 1,000/667 to 128) enabled a shorter length of the descriptor, now compatible with other small molecule and biological descriptors derived in the group^64–66^, and alleviated the overall computational cost of the methodology. An illustrated example of low- and high-entropy lead-like molecules is shown in **Fig 1c**.

Overall, and in view of the benchmark results, we established that the use of rigid docking (rDock) and 128 LLM was the standard and optimal methodology to generate PocketVec descriptors. All results presented along the rest of the manuscript were derived following this strategy. The selected LLM are shown in **Fig S8** and can also be found in our GitLab repository in SMILES and SDF format.

## Results and discussion

### PocketVec performance on the ProSPECCTs benchmark sets

The performance of PocketVec descriptors across ProSPECCTs datasets is assessed in terms of the AUROC and is shown in **Fig 1d**. We observe that PocketVec descriptors are robust to varying definitions of the same pocket, as determined by different crystallized ligands (P1, AUROC 0.97). When restricting such definitions to chemically similar ligands, the performance is maximal (P1.2, AUROC 1.00). Similarly, our descriptors are robust against protein conformational changes, i.e. protein flexibility (P2, AUROC 0.96), and they are also able to distinguish identical pockets from those altered by 5 artificial mutations leading to changes in physicochemical and shape properties of pocket-lining residues. In these cases, we obtained more modest performances (P3 - physicochemical changes and P4 - both physicochemical and shape changes, AUROCs of 0.67 and 0.72, respectively). Reassuringly though, we observed a significant correlation between the number of artificial mutations (1 to 5) and the corresponding AUROC values (Pearson CC > 0.98, p-value < 0.005 in both P3 and P4, **Fig S9**), which confirmed that an increasing number of mutations in the binding site came along with an improved ability to detect such differences using PocketVec descriptors.

We also benchmarked our descriptors in more biologically relevant scenarios where, for instance, pockets binding similar small molecules are found in structurally different proteins. ProSPECCTs includes two datasets to address such cases: P5 includes pocket structures classified into 9 distinct ligand classes (e.g. HEM, ATP, NAD, etc.), and P7 includes a realistic set of binding site pairs reported to be similar in published literature, some of them identified in otherwise unrelated proteins. In these sets, PocketVec descriptors show performances with AUROCs of 0.64 and 0.87, respectively, demonstrating their ability to identify similar pockets in globally dissimilar proteins. It is important to note that two binding sites having identical (or chemically similar) crystallized ligands are not necessarily similar from a PocketVec perspective. Following the logic behind the similarity ensemble approach (SEA^57^), in which targets are quantitatively compared based on the chemical similarity of the ensemble of their ligands, a pair of targets sharing a single active compound may not be significantly similar.

Finally, with the goal of defining a PocketVec distance threshold to classify any pocket pair of interest as either similar or dissimilar, we analyzed the behavior of the Matthew’s Correlation Coefficient (MCC) at multiple cut-off values for the ProSPECCTs datasets (**Fig S10**). In this way, we defined 0.17 as our PocketVec threshold distance, which was the value maximizing the MCC in ProSPECCTs P1.

For the sake of completeness, we generated results for all combinations of docking methods (rigid - rDock, and flexible - SMINA) and reduced sets of chemical compounds (128 LLM and 128 fragments). The use of rDock and LLM (128) was still the best strategy according to the ProSPECCTs datasets (**Fig S11**). Detailed plots for all ProSPECCTs datasets including ROC Curves, PR Curves and distributions of PocketVec distances, docking scores and pocket volumes are included in our GitLab repository (see *Code and data availability*).

### Comparison with existing strategies

Next, we compared the performance of PocketVec descriptors with state-of-the-art methodologies, as reported in^25, 26, 41^, where the authors benchmarked several pocket comparison tools against the ProSPECCTs datasets, including six strategies based on pocket descriptors (**Fig 1d** and **Fig S11**). Overall, PocketVec is the second-best ranked strategy in terms of the weighted average among datasets (0.89) and, indeed, it surpasses the median and the average performance in all ProSPECCTs datasets, apart from P3 (**Table S1**). The top-scoring method is SiteAlign^19^, which is alignment-dependent and based on the projection of residue descriptors into a triangle-discretized sphere that quantifies binding site similarity by minimizing distances between systematically generated cavity fingerprints obtained by moving one binding site with respect to the other. Thus, being specifically developed to compare pockets, it does not provide a unique descriptor for each binding site, which hampers the exploration of the pocket space in the same way molecular fingerprints do for the chemical space of small molecules. Other strategies (e.g.^21, 22, 25, 26^) perform similarly to PocketVec throughout ProSPECCTs datasets, but strongly depend on training data and provide pocket embeddings that are difficult to interpret. Thus, overall, PocketVec represents a fast strategy to generate accurate pocket descriptors that overcome the aforementioned limitations, and it shows a higher performance at assessing pocket similarities than most current strategies (**Fig S12**).

### Comprehensive characterization of druggable pockets in the human proteome

PocketVec descriptors constitute an optimal framework to comprehensively characterize large and diverse sets of small molecule binding sites (apo/holo structures), enabling the navigation across the pocket space of complete proteomes. In view of this, we designed a computational pipeline to generate PocketVec descriptors for all pockets included within human protein domains (**Fig 2a**). Please, see *Online Methods f*or a detailed description of the strategy.

**Fig 2:**
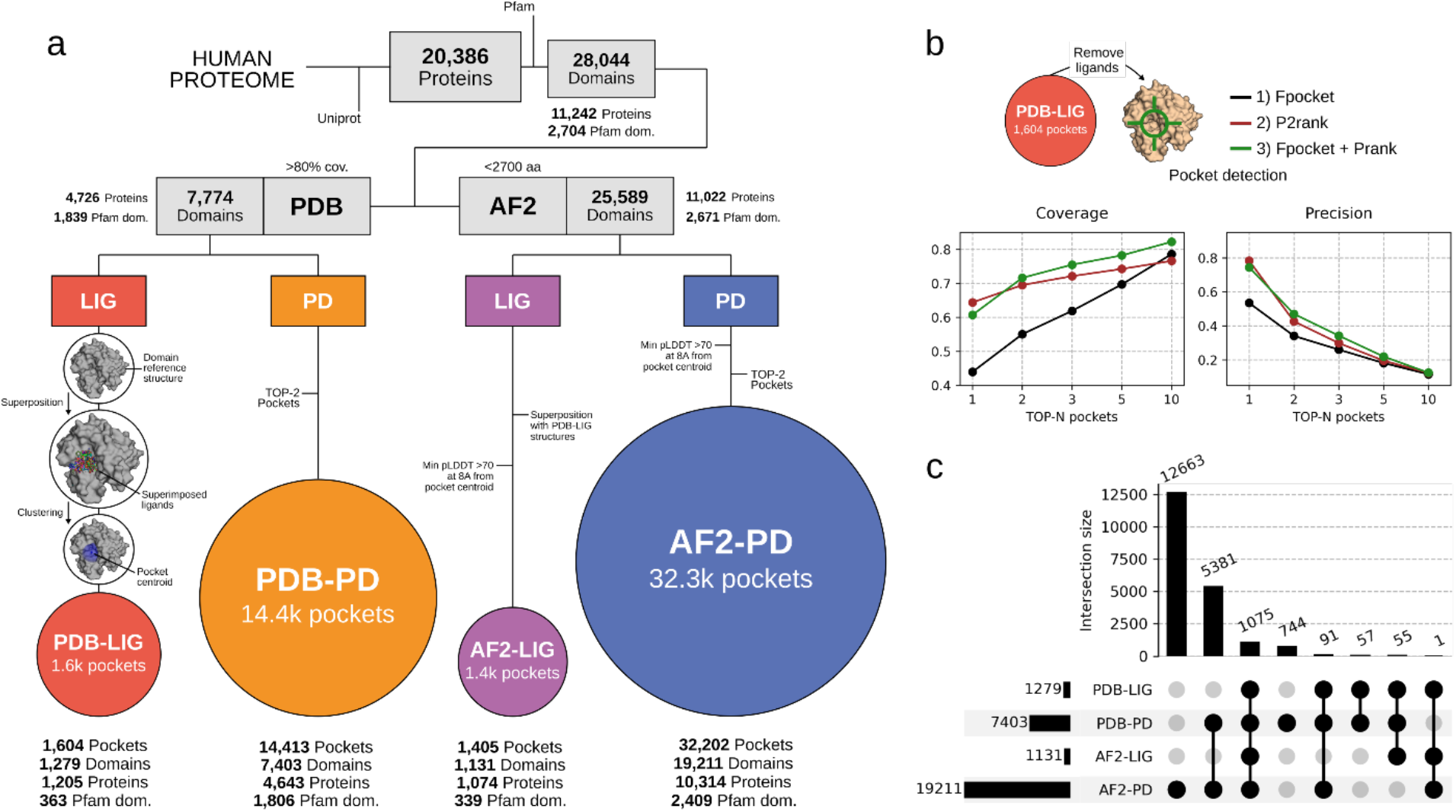
Generating PocketVec descriptors for all druggable pockets within human protein domains. **a)** Computational pipeline to gather all available structural data for human druggable pockets included within protein Pfam domains. Starting from the complete human proteome as per Uniprot, we first identified Pfam domains in experimentally determined structures (PDB) and AF2 predicted models. For PDB structures, we identified pockets in *holo* domain structures based on the position of co-crystallized ligands (1,604 PDB-LIG pockets, red) and in *apo* domain structures using pocket detection techniques (14,413 PDB-PD pockets, orange). We defined ligand-based pockets on AF2 structures (1,405 AF2-LIG pockets, purple) by directly superimposing the location of PDB-LIG pockets, and pockets were also predicted in AF2 models by means of pocket detection algorithms (32,202 AF2-PD pockets, blue). For additional details about the pipeline, please consult the text and the *Online Methods* section. **b)** Benchmark of 3 distinct pocket detection strategies: Fpocket (black), P2rank (brown) and the combination of Fpocket and Prank (green). First, we removed bound ligands from reference PDB structures (PDB-LIG) and we then assessed the performance of the mentioned strategies to detect pockets in ligand-free structures. The plots below represent the evolution of Coverage (proportion of real pockets that are actually detected) and Precision (proportion of detected pockets that are actually real -have a crystallized ligand on PDB-LIG) in terms of the number of best-scoring detected pockets per domain that are considered (1, 2, 3, 5 and 10). **c)** UpSet plot representing the intersecting number of domains among pocket sets (1,279 domains in PDB-LIG, 7,403 domains in PDB-PD, 1,131 domains in AF2-LIG and 19,211 domains in AF2-PD).

In brief, we first retrieved all 20,386 human proteins in UniProt^67^. To avoid working with unstructured or very flexible regions, difficult to model and unlikely to contain druggable pockets, we only kept those protein sequences within autonomous structural units (i.e. domains), as defined in Pfam^68^. Overall, we kept 28,044 domains (2,704 unique Pfam domains) in 11,242 human proteins (**Fig S13**). The next step was to structurally annotate these domains, for which we used two different strategies. On the one hand, we looked for experimentally determined structures searching the PDB^2^, identifying at least one PDB structure for 7,774 domains (1,839 unique Pfam domains in 4,726 proteins). Additionally, we downloaded all human predicted protein structures from AlphaFold DB, obtaining structural models for 25,589 domains (2,671 unique Pfam domains in 11,022 proteins). Indeed, and supporting our strategy, it has been recently shown that the experimental validation rates of small molecule docking is indistinguishable when PDB structures or AF2 models are used^69^.

Next, for each domain, we identified the potentially druggable pockets, also using two different strategies. In the first one, that we termed *ligand-based* pocket definition, we searched for small molecules co-crystallized together with the protein domain, and defined the druggable pocket accordingly. In total, we found at least one PDB structure containing small molecules for 1,279 domains (363 unique Pfam domains in 1,205 proteins). For 503 of these, we only found a single ligand fulfilling our criteria, whereas for 254 of them we could find 10 or more ligands (**Fig S14**). Then, to compile the list of unique ligand-defined pockets, we chose a reference PDB structure for each protein domain, and superimposed all domain structures with the corresponding bound ligands onto the reference. We then used a single-linkage clustering strategy to merge into a single pocket all those ligands whose centroids were at a distance ≤5Å while maintaining the maximum distance between the global centroid of the cluster and the centroids of the individual compounds ≤18Å. We considered the final global cluster centroids as the pocket centroids. Overall, we found 1,604 ligand-defined pockets in 1,279 protein domains (363 unique Pfam domains in 1,205 proteins). We named this set of pockets *PDB-LIG*. To apply the same criteria to those structures modeled with AF2, for each domain, we superimposed the reference PDB structure onto the AF2 model, and transferred the location of the identified PDB-LIG pockets accordingly. We only considered those pockets having a pLDDT value >70 for all residues at a distance ≤8Å from the pocket centroid. In total, we identified 1,405 pockets in 1,131 domains (339 unique Pfam domains in 1,074 proteins), and named this set *AF2-LIG*. As a complementary strategy, and to increase the overall coverage of the human pocketome, we attempted a *de novo* identification of druggable pockets. After testing different methods, we found that the best strategy to detect binding sites in ligand-free structures was the combination of Fpocket^70^, for pocket detection, and Prank^71^ to score them. Using this combination, and considering the top-2 best scored pockets for each domain, we were able to detect 72% of the real pockets while 47% of detected pockets were indeed real (**Fig 2b**, coverage and precision, respectively). Thus, for each domain, we ran Fpocket on the PDB reference structure to identify potential pockets, we ranked them by means of Prank, and we kept the top-2 ranked pockets per domain. Overall, this accounted for a total of 14,413 predicted pockets in 7,403 domains (1,806 unique Pfam domains in 4,643 proteins). We named this set *PDB-PD*. We then used the same strategy and criteria as before to detect pockets onto the predicted AF2 domain models, annotating a total of 32,202 pockets in 19,211 domains (2,409 unique Pfam domains in 10,314 proteins). We named this set *AF2-PD*.

### Robustness in the detection of druggable pockets

Globally, using the different strategies to structurally annotate human protein domains and to identify validated and potential druggable pockets, we compiled 1,604 PDB-LIG, 1,405 AF2-LIG, 14,413 PDB-PD and 32,202 AF2-PD druggable pockets, in 20,067 domains (**Fig 2a** and **Fig 2c**). The significant added value of *de novo* methodologies is very apparent. Indeed, for 18,788 domains (93.6%), all pockets were identified only with pocket detection strategies and, among these, 12,663 domains (63.1%) exclusively featured pockets on AF2 models (**Fig 2c**).

Interestingly, when we assessed the methodological robustness of pocket predictions using a subset of 1,000 PDB-PD domains, we observed that results slightly differed after translating and rotating structures. This effect was observed for both PDB and AF2 structures in a consistent manner: only ∼85% and ∼86% of detected pockets were evenly identified after rotation and translation, respectively (top-2 pockets, **Fig S15a**). However, in the pockets identified regardless of variations in the initial structures, the scoring was very consistent both in PDB structures and AF2 models (Pearson CC ∼0.98). We also evaluated the coherence between pockets detected in PDB and AF2 structures, finding that only ∼59% and ∼49% of detected pockets were evenly identified in PDB and AF2 structures with respect to a reference PDB structure. However, as when comparing discrepancies due to different initial orientations, in the pockets identified both in PDB and AF2 structures the scoring was quite robust, with Pearson CC of ∼0.88 and ∼0.62 for PDB and AF2 models, respectively (**Fig S15b**).

As a final check, we also explored potential differences in the physicochemical properties between the real (ligand-defined) and predicted (detected) pockets, comparing their volumes and buriedness values (see *Online Methods*). In this case, as the only remarkable difference, we found that real pockets identified from bound ligands tended to be smaller than those predicted in the PD framework, with average volumes of ∼3,000 Å^3^ and ∼3,800 Å^3^ for LIG and PD pockets, respectively (**Fig 3a**). As expected, we reached the same conclusion with buriedness values (**Fig S16**), since pocket volume and buriedness were indeed negatively correlated (**Fig S17**).

**Fig 3:**
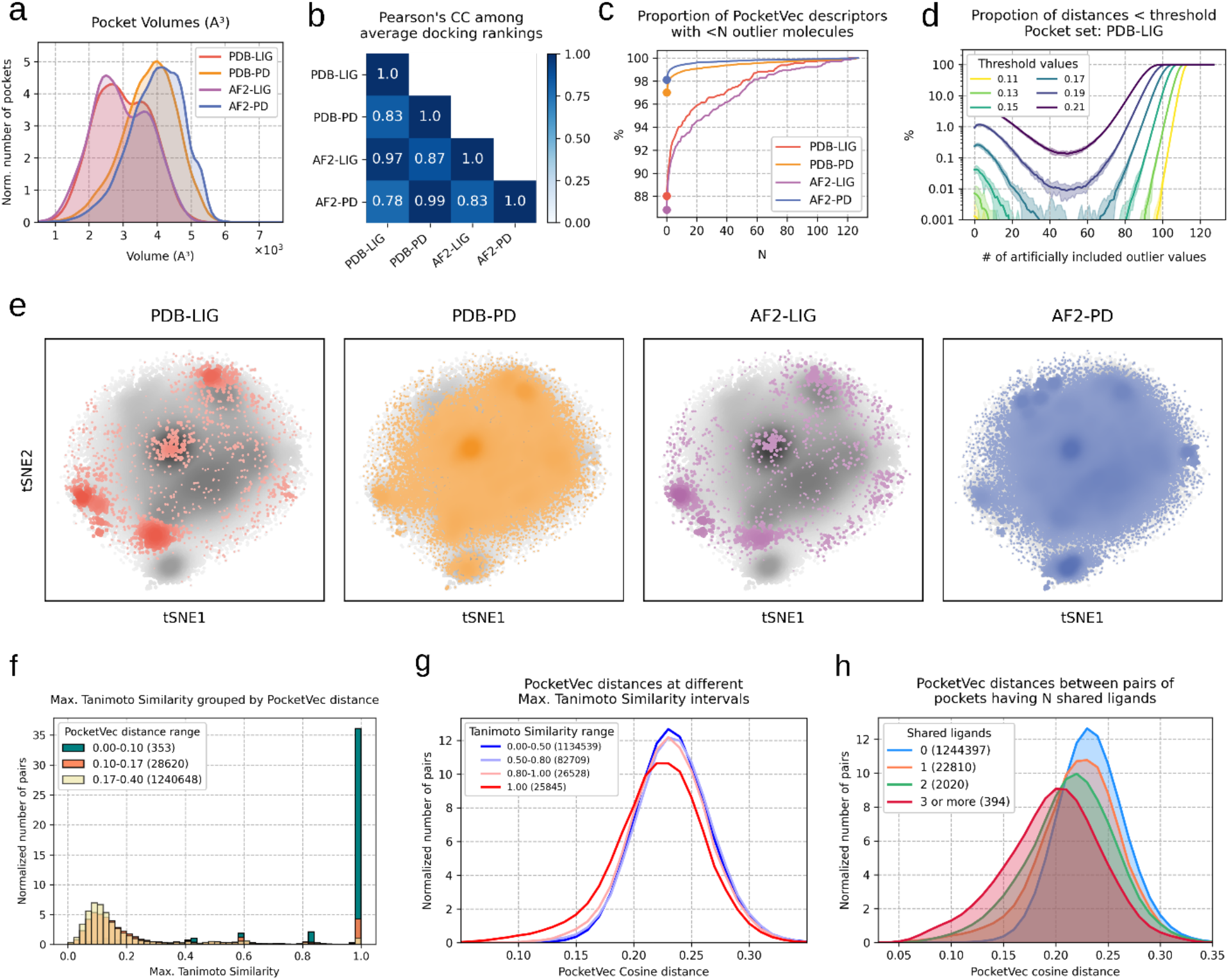
Characterization of human druggable pockets using PocketVec descriptors. **a)** Distribution of volumes (x-axis, x10³ A³) for each pocket set: PDB-LIG (1,604 pockets, red), PDB-PD (14,413 pockets, orange), AF2-LIG (1,405 pockets, purple) and AF2-PD (32,302 pockets, blue). All distributions are normalized (density, y-axis). The average volumes are 2992.88 A³, 3764.1 A³, 2940.47 A³ and 3972.66 A³, respectively. **b)** Correlation between average docking rankings among all pocket sets. For each pocket set, a 128-length array containing the average docking ranking for each lead-like molecule was calculated, and these arrays were compared among pocket sets by means of the Pearson’s Correlation Coefficient. All p-values (10) are <10^-25^. **c)** For each pocket set, proportion of PocketVec descriptors (%, y-axis) having less than N (x-axis) outlier molecules (i.e. molecules with positive docking scores, please see *Online Methods*). **d)** Proportion of originally dissimilar pocket pairs (y-axis, distances >threshold) classified as similar (distances <threshold) due to the artificial insertion of a growing number of outlier values (x-axis) at different distance thresholds. To run the analysis, 100,000 PocketVec distances were randomly sampled (x10 times) from PDB-LIG. Solid lines represent average results and shadowed areas correspond to minimum and maximum proportions among samples. Results did not change among pocket sets (**Fig S21**). **e)** tSNE (t-distributed Stochastic Neighbor Embedding) representation of all PocketVec descriptors within each pocket set (PDB-LIG, PDB-PD, AF2-LIG, AF2-PD). Each pocket is represented by a single point, which is colored and sized by 2D density within the pocket set. Gray dots correspond to the background space: all PocketVec descriptors are considered at once, and are also colored and sized by 2D density. **f)** Distributions (density, y-axis) of maximum Tanimoto Similarity among bound ligands (x-axis, bin width: 0.02) grouped by PocketVec distance ranges (0-0.10, 0.10-0.17 and 0.17-0.4). The number of pocket pairs per PocketVec distance range is specified in parenthesis. **g)** Distributions (density, y-axis) of PocketVec cosine distances (x-axis) grouped by the maximum Tanimoto Similarity among bound ligands (0-0.5, 0.5-0.8, 0.8-1 and 1). The number of pocket pairs per maximum Tanimoto Similarity range is specified in parenthesis. **h)** Distributions (density, y-axis) of PocketVec cosine distances (x-axis) grouped by the number of shared ligands between pockets (0, 1, 2 and 3 or more). The number of pocket pairs per number of shared ligands is specified in parenthesis.

Altogether, these results underscore some limitations in the pocket detection strategy, revealing slight variations due to the orientation of the initial structures, a stronger dependence on structural variability and the production of predicted pockets whose physicochemical properties (e.g. volume) diverge from known pockets.

### Systematic generation of PocketVec descriptors

Once we had identified human druggable pockets with four different approaches, we systematically generated PocketVec descriptors for all of them, using the strategy described above (**Fig 1a**). The high number of pockets explored enabled an exhaustive analysis of potential dependencies between the lead-like molecules used to build the descriptors and the characterization of druggable pockets. Reassuringly, we did not find any correlation between docking rankings and molecular properties of docked compounds (e.g. molecular weight or number of heavy atoms, **Fig S18** and **Fig S19**, respectively). On the contrary, we observed that most molecules exhibited a complete range of rankings (from 1 to 128), although the ranking distributions showed that some molecules tended to bind with good scores in many pockets while others were mostly downranked (**Fig S20**). Indeed, this tendency was observed in all pocket definition strategies, and we found significant correlations between the average docking rankings for docked lead-like molecules among different pocket sets (**Fig 3b**). Such correlations showed a perfect agreement between PocketVec descriptors generated through PDB and AF2 structures, but exposed slight differences between LIG and PD results (**Fig 3b**, Pearson’s CC <0.88). In line with this, we found that 88.0% and 86.8% of the PocketVec descriptors generated for PDB-LIG and AF2-LIG, respectively, showed no lead-like molecules with positive docking scores (i.e. outlier molecules not fitting in the pocket, see *Online Methods*), while the fraction was considerably higher in PDB-PD and AF2-PD sets, with 97.0% and 98.1% of the descriptors, respectively (**Fig 3c**). This result was consistent with the differences observed in pocket volumes (**Fig 3a**). Reassuringly, we found that smaller pockets (<3,000Å^3^) tended to be more prone to exhibit outlier molecules than bigger pockets (Fisher exact test, OR >70 and p-value <10^-45^ for all pocket sets).

To assess if outlier compounds could have a significant effect in the generation of poor PocketVec descriptors, we randomly inserted outlier values in the descriptors and computed the fraction of originally dissimilar pocket pairs (PocketVec distance >0.17) that were incorrectly labeled as similar (distance <0.17) due to an increasing number of outlier values. We observed that the insertion of up to 80 outlier molecules (out of 128) did not significantly compromise PocketVec descriptors, leading to only ∼0.039% of false positives (**Fig 3d** and **Fig S21**). In front of this strong robustness of the descriptors, in further analyses, we only discarded those very few PocketVec descriptors with more than 80 outlier molecules (10, 45, 15 and 43 descriptors from the PDB-LIG, PDB-PD, AF2-LIG and AF2-PD sets, respectively).

### All-vs-all comparison of human druggable pockets

The vector-like nature of PocketVec descriptors makes them perfectly suited for extremely fast comparisons by computing simple cosine distances, enabling comprehensive proteome-wide similarity searches. We first generated a t-distributed Stochastic Neighbor Embedding (t-SNE) representation for all PocketVec descriptors to facilitate a qualitative visualization of the characterized pocket space (**Fig 3e**). This visualization underlined several interesting points, some of which have already been discussed in previous sections: (i) pockets defined by bound ligands (PDB-LIG and AF2-LIG) predominantly occupy well defined regions of the pocket space, (ii) pocket detection strategies (PDB-PD and AF2-PD) significantly contribute to expand the overall coverage of the human druggable pocket space, (iii) the use of AF2 models particularly bolsters this expansion. Perhaps more interestingly, the PDB-PD and AF2-PD maps highlight dense areas of the human pocket space for which we do not yet have any experimental structure with a bound chemical compound.

Then, we systematically evaluated the validity of the chemogenomic hypothesis behind the generation of PocketVec descriptors (i.e. similar pockets bind similar ligands). We compared all PDB-LIG pockets having PocketVec descriptors (1,594 pockets, ∼1.27M comparisons) on the basis of the maximum Tanimoto similarity among their bound ligands using ECFPs (2,048 bits and radius 2), and we grouped pocket pairs according to their cosine PocketVec distance (**Fig 3f**). We found that, indeed, pocket pairs with small PocketVec distances (<0.10) typically showed higher maximum Tanimoto similarities (>0.85) among their ligands than pocket pairs with high PocketVec distance (>0.17, Fisher’s exact test OR >90 and p-value <10^-300^), thus supporting our underlying hypothesis. However, the fact that similar pockets bind similar ligands does not necessarily imply that similar ligands bind similar pockets. Indeed, we observed that, in general, PocketVec distances did not decrease substantially when increasing the maximum Tanimoto similarity among bound ligands (**Fig 3g**). The only exception was for almost identical ligands (Tanimoto similarities=1), which showed a very subtle deviation towards smaller PocketVec distances (AUROC 0.59, true positives having Max. Tanimoto Similarity=1, true negatives having Max. Tanimoto Similarity in the [0, 0.5) range), in line with the behavior observed in ProSPECCTs P5 (AUROC=0.64) and P5.2 (AUROC=0.62). While this may seem somewhat counterintuitive, it serves as evidence to decipher the quantification of pocket similarity using PocketVec descriptors.

The fact that a single compound is consistently upranked for two pockets may indeed be a first evidence of pocket similarity but falls short to provide similar PocketVec descriptors if no other compound is ranked in a systematic manner. In practical terms, this translates into pockets that share a growing number of ligands being more likely to be similar from a PocketVec perspective, as highlighted by Shoichet and co-workers when they presented their similarity ensemble approach (SEA) to unveil protein remote relationships^57^. Reassuringly, we found that pockets that shared ligands in the PDB-LIG set tended to be more similar (i.e. smaller PocketVec distances) than pockets sharing no ligands (**Fig 3h**). However, the deviation towards smaller distances for pockets sharing a single ligand was subtle (AUROC = 0.58, true positives sharing 1 ligand and true negatives sharing no ligands), but when 3 or more ligands were shared between pockets, the effect was already notable (AUROC = 0.75, true positives sharing 3 or more ligands and true negatives sharing no ligands). We also found that only 2.1% of pocket pairs sharing no ligands in the PDB-LIG set showed PocketVec distances <0.17, while this fraction increased to 26.1% when 3 or more ligands were shared between pockets. However, it is obvious that the lack of shared co-crystallized compounds between two pockets does not necessarily mean that they might not have common ligands. To overcome the limitation of potentially missing ligands, we used a pure computational strategy: we computed docking scores for the 128 standard lead-like molecules (see *Methodological development and implementation*) against all PDB-LIG defined pockets and labeled them as active (the top 1% of docking scores) or inactive (**Fig S22a**). We observed that the vast majority of the lead-like molecules (127 out of 128) were cataloged as active in, at least, one pocket (**Fig S22b**), and almost 20,000 of the potential ∼1,29M PDB-LIG pocket pairs shared, at least, one active lead-like molecule (**Fig S22c**). With this alternative approach, we confirmed the tendency observed using experimental data: the more shared ligands between pockets, the smaller their PocketVec distance (AUROC = 0.87 when comparing pockets sharing no ligands and those sharing 3 or more ligands; **Fig S22d**).

Next, we explored the complementarity and added value of PocketVec descriptors with respect to more established strategies to compare protein families and druggable pockets, such as sequence and structure similarity. First, we sought to investigate the correlation between sequence, structure and PocketVec similarity (defined as 1-PocketVec distance) among pockets located in the same Pfam domains in the more comprehensive set of AF2-PD pockets (see *Online Methods*). As expected, we found that the higher the sequential and structural similarity between compared pockets (assessed by sequence identity and Cα RMSD, respectively), the more similar they were according to PocketVec descriptors (Pearson CC of 0.55 and -0.35, respectively, p-value<10^-100^ in both cases; **Fig 4a**). Reassuringly, the observed correlations were also found when computed on AF2-LIG pockets (Pearson CC of 0.57 and -0.35 for sequence identity and Cα RMSD, respectively **Fig S23**). However, there were indeed cases where the results did not align perfectly, underscoring the distinctive and complementary insights that PocketVec descriptors can provide beyond traditional sequential and structural analyses.

**Fig 4:**
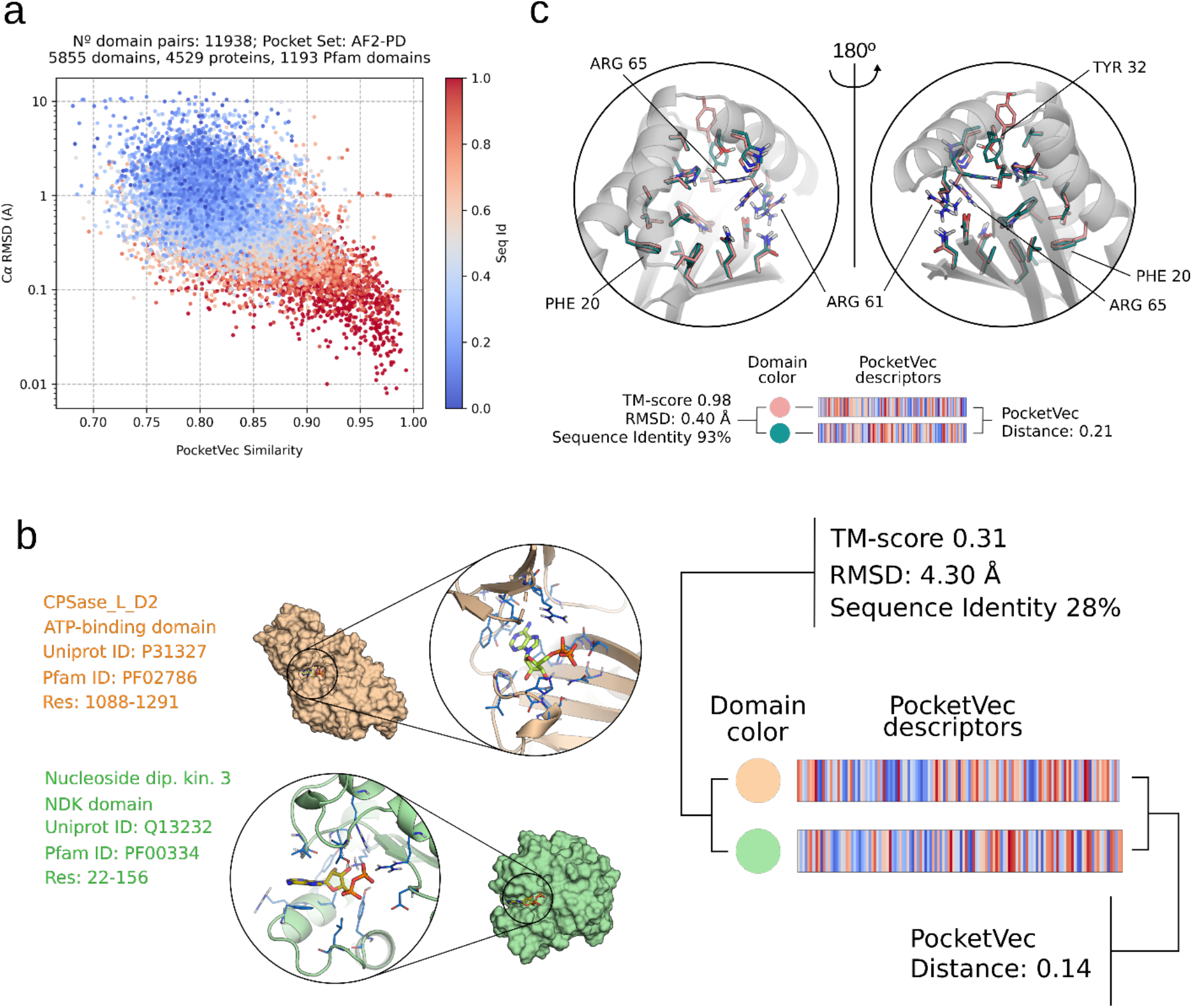
Using PocketVec descriptors to assess proteome-wide pocket similarity: achieving otherwise unattainable insights. **a)** Correlation between PocketVec similarity (x-axis, defined as 1-PocketVec distance), structural similarity (y-axis, Cα RMSD) and sequence identity (color) among pockets located at the same Pfam domains (max. 10) in the AF2-PD pocket set. Pearson CC between PocketVec similarity and sequence identity: 0.55 (p-value∼0). Pearson CC between PocketVec similarity and RMSD: -0.35 (p-value∼0). **b)** Similar pockets found in dissimilar domains. Pockets were found in the CPSase_L_D2 ATP-binding domain (PF02786, top structure, wheat color) of the Carbamoyl-phosphate synthase (P31327, positions 1088-1291. PDB ID: 5DOU) and in the NDK domain (PF00334, bottom structure, pale green color) of the Nucleoside diphosphate kinase 3 (Q13232, positions 22-156. PDB ID: 1ZS6). Pockets (both in the PDB-LIG set) have a PocketVec distance of 0.14 (below the established threshold of 0.17, please see *PocketVec performance on the ProSPECCTs benchmark sets*) although domains share a sequence identity of only 28% and a poor structural similarity (TM-score=0.31, RMSD=4.3Å). **c)** Dissimilar pockets found in similar domains. Pockets were found in the NIPSNAP domain (PF07978) of the Protein NipSnap homolog 3B (Q9BS92, positions 146-245, AF2 model, pink residues) and in the NIPSNAP domain (PF07978) of the Protein NipSnap homolog 3A (Q9UFN0, positions 146-245, AF2 model, green residues). The former is used as the reference structure (gray cartoon). Pockets (both in the AF2-PD set) have a PocketVec distance of 0.21 (above the established threshold of 0.17, please see *PocketVec performance on the ProSPECCTs benchmark sets*) although domains have a sequence identity of 93% and also a very high level of structural similarity (TM-score=0.98 and RMSD=0.4Å).

Additionally, we also ran an all-against-all pocket comparison within and across pocket sets (PDB-LIG, PDB-PD, AF2-LIG and AF2-PD), computing over 1.2 billion pocket comparisons. Interestingly, we found more than 3.5 million similar pockets in domains having low sequential and structural similarities (PocketVec distance <0.17; TM-score <0.35; Sequence identity <30%). For instance, we found similar pockets (PocketVec distance: 0.14, both pockets in the PDB-LIG set) in the CPSase_L_D2 ATP-binding domain (PF02786) of the Carbamoyl-phosphate synthase (P31327, positions 1088-1291) and in the NDK domain of the Nucleoside diphosphate kinase 3 (Q13232, positions 22-156), although they shared a sequence identity of only 28% and a poor structural similarity (TM-score=0.31, RMSD=4.3Å). Reassuringly, crystal structures confirmed that both pockets can bind ADP (PDB IDs: 5DOU and 1ZS6, respectively), which strengthened our observation that these pockets were indeed similar (**Fig 4b**). The inventory of all similar pockets (PocketVec distance <0.17) together with structural and sequential comparisons at domain level are reported in our GitLab. On the other hand, our analyses also revealed more than 29k pocket pairs (out of a subsample of 11.1 million pairs having PocketVec distance >0.20) that, despite being similar in terms of sequence and structure, showed quite dissimilar pockets (PocketVec distance >0.20; TM-score >0.50; Sequence identity >40%). As an illustrative example, we identified different druggable pockets (PocketVec distance: 0.21, both pockets in the AF2-PD set) in the NIPSNAP domain (PF07978) of the Protein NipSnap homolog 3B (Q9BS92, positions 146-245) and in the NIPSNAP domain (PF07978) of the Protein NipSnap homolog 3A (Q9UFN0, positions 146-245), although these domains had a sequence identity of 93% and also a very high level of structural similarity (**Fig 4c**, TM-score=0.98 and RMSD=0.4Å).

### Identifying kinases with similar inhibition profiles through PocketVec descriptors

Protein kinases have long been a cornerstone of drug discovery efforts primarily due to their role as oncogenic targets^72^. Since the breakthrough FDA-approval of imatinib in 2001, dozens of small molecules (72 as of November 2022^73^) have been approved as therapeutic kinase inhibitors for clinical use. However, the design of selective inhibitors is a challenging task due to the highly conserved ATP-binding pocket shared among human kinases. Indeed, many kinase inhibitors show high promiscuity (i.e. they bind to many kinases), while others are rather selective^74, 75^. This same variability is also apparent from the kinase perspective: some bind to many inhibitors while others are extremely selective^76^. Our characterization of druggable pockets in human proteins showed 1,286 potential binding sites within protein kinase domains. More specifically, we derived PocketVec descriptors for 229, 404, 195 and 458 pockets in the PDB-LIG, PDB-PD, AF2-LIG and AF2-PD sets, respectively. Thus, we set up to explore a potential correlation between the pockets in the different kinases and their experimentally determined small molecule inhibition profiles.

We first collected kinase-inhibitor pairs identified through systematic chemical proteomics approaches by Kuester and collaborators, where they tested interactions between 520 kinases and 243 inhibitors (2017 set)^77^, and between 318 kinases and 1,183 inhibitors (2023 set)^78^. We used a standard activity cutoff of 30nM to define binary kinase-inhibitor matrices, as recommended in Pharos 17 (http://pharos.nih.gov). In the 2017 set, we found interactions involving 111 kinases and 94 inhibitors (**Fig 5a**), with 43 kinases being inhibited by a single compound, and 18 of them interacting with 5 or more inhibitors (**Fig S24a**). In the 2023 set, we identified interactions comprising 73 kinases and 164 inhibitors, where 19 kinases interacted with 1 inhibitor and 27 with 5 or more (**Fig S24b**). We then compared kinases on the basis of their binarized inhibition profiles (Jaccard similarity, **Fig 5d, j**, upper triangular matrices), and we observed that a relatively small fraction of kinase pairs shared at least 1 inhibitor (12.3% and 9.7% for the 2017 and 2023 sets, respectively). We also compared the kinases using PocketVec descriptors (**Fig 5d, j**, lower triangular matrices), finding that the fraction of kinase pairs showing similar druggable pockets (i.e. PocketVec distances <0.17) was 40.5% and 41.4%, respectively. As expected, these fractions were significantly higher than the figure obtained when comparing random sets of pocket descriptors (∼2%. Fisher’s exact test, OR >30, p-value ∼ 0), since all kinases contain at least one similar ATP-binding pocket. Reassuringly, as observed in the general analysis of human druggable pockets (**Fig 3h**), we found that the higher the number of common inhibitors between pairs of kinases, the more similar their pockets were according to PocketVec descriptors (**Fig 5b, h**). However, even when no shared inhibitor was found for a pair of kinases, as expected, the similarity between their ATP-binding pockets was quite remarkable, suggesting that the PocketVec distance threshold should be lowered to disentangle drug promiscuity.

**Fig 5:**
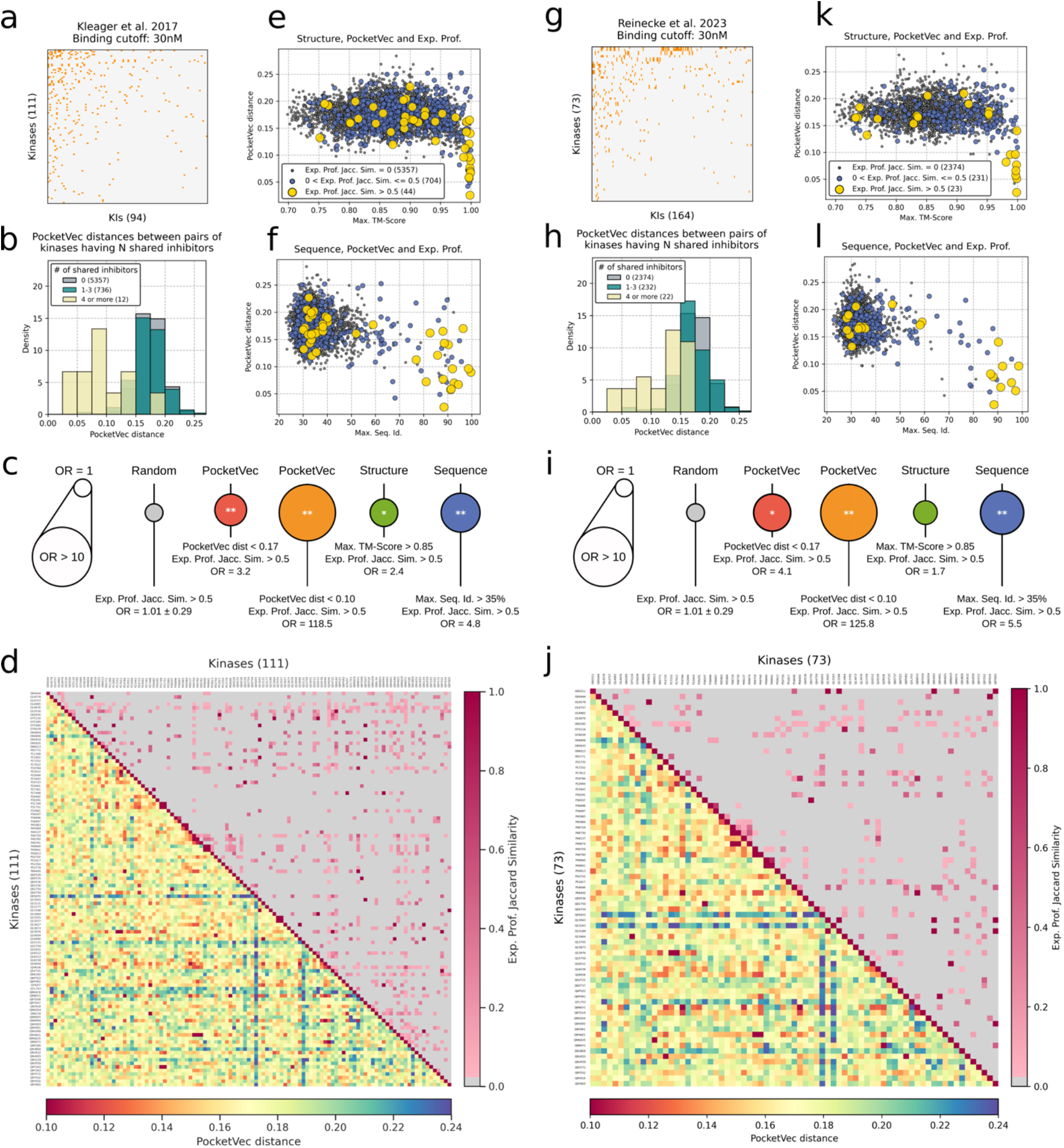
Correlation between inhibition profiles and PocketVec descriptors. All the analyses have been performed on the data obtained from Kleager et al.^77^ (right panels) and Reinecke et al.^78^ (left panels). **a, g)** Inhibition matrix between protein kinases, and small molecule kinase inhibitors binarized at 30 nM. Both kinases and inhibitors are sorted by the number of active inhibition events. Orange dots indicate inhibition and white dots indicate no inhibition. **b, h)** Distributions of PocketVec distances grouped by the number of shared inhibitors between kinases (0, 1-3 and 4 or more). The number of kinase pairs per number of shared inhibitors is specified in parenthesis. **c, i)** Enrichments (Fisher’s exact test) in similar inhibition profiles (Jaccard Similarity >0.5) for those kinase pairs being similar in terms of PocketVec distance (red: <0.17, orange: <0.10), structural similarity (TM-score >0.85) and sequence identity (>35%). For comparison, the results obtained with randomly selected kinase pairs (gray) are also included. Circle areas are proportional to the corresponding ORs and p-values are specified in the center with the following format: * p-value < 0.05, ** p-value < 0.001. **d, j)** Pairwise kinase comparisons. Rows and columns correspond to alphabetically sorted kinases (by Uniprot ID). Upper triangular matrices: kinases are compared on the basis of their experimentally determined inhibition profiles. Each square represents the Jaccard similarity between the inhibition profiles of two targets: the higher the Jaccard similarity, the more similar the corresponding inhibition profiles. Lower triangular matrices: kinases are compared on the basis of their PocketVec descriptors (employing the minimum distance among all PocketVec descriptors in the Protein Kinase Domains). The color of each square indicates the minimum PocketVec distance between two targets: the lower the PocketVec distance (red), the more similar the kinases are at pocket level according to our descriptors. **e, k)** Relationship between structural similarity (x-axis, Max. TM-score) and PocketVec distances (y-axis) between pairs of protein kinases. Each point represents a kinase pair and is colored and sized in terms of the similarity between experimentally determined inhibition profiles. **f, l)** Relationship between sequence similarity (x-axis, Max. Seq. Id.) and PocketVec distances (y-axis) between pairs of protein kinases. Each point represents a kinase pair and is colored and sized in terms of the similarity between experimentally determined inhibition profiles.

Next, we parsed the general inhibition matrices to derive inhibition profiles for each kinase (i.e. a vector describing whether it does or does not interact with every tested inhibitor), and we assessed the coherence between PocketVec distances and the similarity between experimentally determined inhibition profiles. Indeed, kinase pairs with low PocketVec distances (<0.17) showed a significant enrichment for similar inhibition profiles (Fisher’s exact test, OR = 3.2, p-value <0.0003 in the 2017 set, and OR = 4.1, p-value <0.003 in the 2023 set for a Jaccard similarity >0.5). These enrichments were even more pronounced when we applied more stringent distance thresholds (PocketVec distance <0.10) with OR >118 and OR >125 (p-values <10^-10^) for the 2017 and 2023 sets, respectively (**Fig 5c, i**).

We then compared our results using PocketVec descriptors to more classical structure and sequence similarity analyses (see *Online Methods*). As expected, we found that structurally similar proteins (Max. TM-score >0.85) were also moderately enriched in similar inhibition profiles (OR = 2.4, p-value <0.05; OR = 1.7, p-value >0.05), as were kinases sharing >35% sequence identity (OR = 4.8, p-value <10^-6^; OR = 5.5, p-value <10^-4^). Finally, we studied the relationship between structure, sequence, PocketVec and inhibition profile similarity between kinase pairs. Consistently with our global analysis of the human druggable pockets (**Fig 4a**), higher structure and sequence similarities consistently led to lower PocketVec distances in both the 2017 (**Fig 5e, f**) and the 2023 (**Fig 5k, l**) sets.

Despite the coherence of the results of the three metrics, interestingly, we found 9 kinase pairs (4 in the 2017 and 5 in the 2023 sets) with clear inhibition profile similarities that only PocketVec descriptors could pick. On the other hand, our analyses also revealed 8 cases (5 in the 2017 and 3 in the 2023 sets) where PocketVec descriptors fell short to detect similarities that could be recovered by sequence or structure comparisons alone. Overall, these results emphasize the coherence and complementarity between strategies, highlighting the potential of PocketVec descriptors to identify otherwise unattainable similarities in experimental inhibition profiles among protein kinases.

### Code and data availability

All the code necessary to generate PocketVec descriptors for any pocket of interest is available in our GitLab (https://gitlabsbnb.irbbarcelona.org/acomajuncosa/pocketvec), together with installation instructions and guidelines for running the code. On average, the generation of a PocketVec descriptor for a protein pocket takes 1h on an Intel Xeon Gold 6138 CPU.

In addition, in the same repository under the *results* directory, we also provide all the pre-computed descriptors and the comparisons with sequence and structure similarity measurements, as described in the text.

## Concluding remarks

We have presented PocketVec, a novel approach to generate vector-like protein pocket descriptors based on inverse docking and the chemogenomics principle that similar pockets bind similar ligands. A thorough assessment of its performance ranks it among the best available methodologies to characterize protein druggable pockets, while overcoming some important limitations. We have also systematically searched for druggable pockets in the folded human proteome, using experimentally determined protein structures and AF2 models, identifying over 32,000 binding sites in more than 20,000 protein domains. We then derived PocketVec descriptors for each small molecule binding site and took advantage of their vector-like format to run an all-against-all pockets similarity search, exploring over 1.2 billion pairwise comparisons. We found that PocketVec descriptors are complementary to other, more classical, search strategies, enabling the identification of pocket similarities not revealed by structure- or sequence-based comparisons. Indeed, a systematic comparison of druggable pockets in protein kinases showed that kinase pairs with similar PocketVec descriptors also exhibited similar experimentally determined inhibition profiles.

This first generation of descriptors has been primarily designed for global analyses, such as the comprehensive characterization of all human druggable pockets. Indeed, our analyses have revealed dense clusters of similar pockets in distinct proteins for which no inhibitor has yet been co-crystalized, opening the door to strategies to prioritize the development of chemical probes to cover the druggable space^79^. Moreover, our initial descriptors can be easily adapted to cater to specific tasks (i.e. exploring substrate specificity in a given protein family) by refining the selection of predefined lead-like molecules used or fine-tuning the similarity cutoff, thereby enhancing their performance. Of special interest are the anticipation of undesired off-targets as well as the guidance of rational polypharmacology, where single univalent molecules could be designed to target two proteins simultaneously, provided that their druggable pockets are similar enough^80^. However, the main impact is likely to come from proteochemometric approaches, where a combination of ligand and target descriptors are used to train machine learning models^13^. It has been shown that structure-based descriptors of the targets are often superior to distinguish drug selectivity, although the sequence-based ones are often used when key protein structural details are lacking^39^. The generation of accurate descriptors derived for not yet described pockets in AF2 protein models overpasses this limitation, and open up many possibilities. We envisage a scenario where small molecule and pocket descriptors combined are used to train AI-based generative models (e.g.^81, 82^ to design new chemical entities that bind each protein druggable cavity. Indeed, the estimated space of 10^33^ synthetically accessible drug-like molecules is mostly unexplored and represents a reservoir of potentially bioactive compounds^83^. Deep learning strategies have successfully designed new antibiotic scaffolds^84^ and placed 15 AI-designed drugs in clinical trials, including first-in-class molecules against several targets^85^. Overall, accurate descriptors of druggable pockets might serve as a cornerstone for the development of generative AI approaches in drug discovery, offering unprecedented opportunities to expedite the design of a chemical toolbox to probe biology and, ultimately, to new therapeutics.

## Online Methods

### Selection of compound sets

Fragments: we downloaded the Glide^60, 61^ diverse fragment dataset from the Schrodinger website (https://www.schrodinger.com) in November 2020. This collection of compounds is composed of 667 molecules having molecular weights the 50-200 g·mol^-1^ range (**Fig S2**).

Lead-like molecules (LLM): we retrieved a set of 650k lead-like molecules from MOE v2019.01 (Chemical Computing Group, Montreal, Canada). We then performed a k-means clustering using TAT fingerprints, setting the number of clusters to 1,000, and selected the corresponding 1,000 molecules closest to each cluster centroid to build the library. These compounds exhibit molecular weights in the 200-450 g·mol^-1^ range (**Fig S2**).

### Small molecule docking strategies

Rigid docking: we performed rigid docking calculations using rDock^62^ (downloaded on July 2021 from https://github.com/CBDD/rDock). We prepared the protein structures using MOE v2019.01 (Chemical Computing Group, Montreal, Canada), Biopython^86^ and the structure checking utility from BioExcel Building Blocks^87^. We ran all docking calculations using standard parameters and scoring functions. The binding site box was built around the pocket centroid (ligand centroid or detected pocket centroid) with a radius of 12 Å (*RbtLigandSiteMapper* option) and the number of runs was set to 25. Finally, we set the *DIHEDRAL_MODE* to *FIXED* (rigid docking). The considered score for each molecule was the minimum value of SCORE.INTER.

Flexible docking: we used SMINA^63^ (downloaded on November 2020 from https://sourceforge.net/projects/smina/) for the flexible docking calculations. We prepared the protein structures using Reduce^88^, OpenBabel^89^, Biopython^86^ and the structure checking utility from BioExcel Building Blocks^87^. We ran all the calculations using standard parameters and scoring functions for flexible docking. The binding site box was automatically derived from the position of the bound ligand (*autobox_ligand* parameter). The considered score for each molecule was the minimum value of *minimized_affinity*.

### Post-docking analysis

For each pocket under evaluation, docking scores were stored in a one-dimensional NumPy array^90^ and then translated into rankings using SciPy^91^ (*rankdata* function, method *max*). In this way, the molecule with the lowest docking score was assigned the top ranking (1^st^), while the one with the highest docking score was ranked as the N^th^ (being N the total number of tested molecules; N=128 in the standard PocketVec pipeline). It is important to note that molecules yielding positive docking scores (e.g. due to structural clashes with the protein) were not explicitly considered and their corresponding rankings were set to an outlier value (e.g. 129). The rationale behind this procedure was that such outlier molecules were indeed informative (i.e. the pocket was too small to fit them) but needed to be distinguished from binders having poor (but negative) docking scores (i.e. weak binders). Specifically, docking scores in the range 0-50, 50-100 and >100 were translated into N+1, N+2 and N+3 rankings, respectively (129, 130, 131 in the standard PocketVec pipeline).

### Benchmark set

A good strategy to identify the best combination of small molecules and docking methods to develop pocket descriptors, and to assess their validity, is to see if they can faithfully capture reported similarities between small molecule binding pockets. To this end, we used ProSPECCTs^41^, a collection of 10 datasets composed of protein-ligand binding site pairs classified as similar or dissimilar according to specific criteria (downloaded in July 2021 from http://www.ewit.ccb.tu-dortmund.de/ag-koch/prospeccts/). P1 (P1.2) includes 326 (45) protein-ligand complexes involving 12 (12) different proteins, and it is meant to study the sensitivity of pocket comparison tools to the binding site definition by comparing proteins having identical sequences with chemically distinct (similar) ligands located at the same site. P2 comprises 17 PDB files resolved by NMRs, containing a total of 329 different models, and was designed to assess the impact of protein flexibility in pocket comparisons. P3 and P4 include a variable number (1 to 5) of randomly added artificial mutations in the P1 proteins leading to changes in the physicochemical (P3) and physicochemical and shape (P4) properties of the protein binding site. For the sake of simplicity and coherence with reported performances, we have only considered structures with 5 mutations (representing 326 out of 1630 mutated structures). P5 (P5.2) was designed to detect pairs of unrelated proteins binding to identical or similar ligands, and consists of 80 (100, including phosphate binding sites) protein-ligand complexes^92^. P6 (P6.2) is intended to evaluate the identification of distant relationships between pockets binding to identical ligands but having variable pocket environments^93^, and it includes 115 protein structures excluding (including) cofactors. We did not use P6 or P6.2 to benchmark our methodology since all pocket pairs are bound to identical or highly similar ligands, and the similar/dissimilar classification is done considering fine details of protein-ligand binding, such as the involved ligand functional groups. Thus, this set is not appropriate to guide and assess the developments of our pocket descriptors. Finally, P7 was retrieved from published successful applications of pocket similarity studies in a diverse set of proteins, and it contains 1,151 protein structures. A detailed overview of all ProSPECCTs datasets is presented in **Fig S3**.

### Entropy measurements

For each molecule within each ProSPECCTs dataset, rankings were first binned into 100 different groups (bins) in order to discretize a variable (rankings) that, in practical terms, was continuous (since ranking range was usually higher than the number of considered structures). Shannon’s Entropy was then calculated using such binned data (SciPy^91^, *entropy* function, base 2).

### Domain-based characterization of the human druggable pockets

We searched all human protein identifiers from UniProt (July 2022, *organism_id 9606* and *reviewed* set to *true*), retrieving a total of 20,386 unique human proteins^67^. Then, for each human protein, we retrieved all Pfam domains^68^, considering only those entities labeled as ‘domain’ (e.g. we did not include ‘repeats’). Overall, we found 28,044 domains (2,704 unique Pfam domains) in 11,242 human proteins (**Fig 2a** and **Fig S13**).

To structurally annotate these domains, we used two different strategies. On the one hand, we looked for experimentally determined structures searching the PDB^2^. For each human domain, we gathered all PDB chains showing a structural coverage of the domain ≥80% using the localpdb package^94^ (PDB version 2022.02.25). We identified at least one PDB structure for 7,774 domains (1,839 unique Pfam domains in 4,726 proteins), processing all PDB files and removing those regions outside the domains under study. Additionally, we downloaded all predicted structures for Homo Sapiens from AlphaFold DB (https://alphafold.ebi.ac.uk/download#proteomes-section, proteins having <2,700 amino acids, August 2022), processed all files and removed those regions that did not match the domains under study, leaving predicted structures for 25,589 domains (2,671 unique Pfam domains in 11,022 proteins).

To identify druggable pockets, we also followed two complementary strategies.

#### Ligand-based

In the ligand-based pocket definition, we identified all the PDB structures (chains) corresponding to human protein domains that contained small molecules co-crystallized with the domains of interest that fulfilled the following criteria: i) Number of carbon atoms >6, to filter out solvent molecules and crystallography-related species and ii) solvent accessibility ≤0.4 or buriedness ≥15. We defined solvent accessibility as the ratio between the ligand solvent-accessible surface area (SASA) in the bound state and the ligand SASA in the free state. SASA values were calculated with RDKit. Additionally, we defined buriedness as the number of protein residues having a distance below 8Å to the ligand centroid, and we calculated these values using Biopython^86^. The cut-off values for accessibility (0.4) and buriedness (15) were set upon visual inspection of many bound ligands. We considered both parameters in order to recover as many interesting cases as possible: accessibility values usually underestimated pockets defined by small ligands while buriedness values often underrated large pockets. In total, we found at least one PDB structure containing a ligand fulfilling all conditions for 1,279 domains (363 unique Pfam domains in 1,205 proteins). For 503 of these, we only found a single ligand, whereas for 254 of them we could find 10 or more ligands (**Fig S14**), including a variety of metabolic nucleotides such as ADP (found in 114 domains) or GDP (found in 100 domains).

To compile the list of unique ligand-defined pockets we followed the procedure shown in **Fig 2a**. First, we chose a reference PDB structure for each protein domain where we considered the structural coverage of the domain and the resolution of the crystal structure. We then superimposed all domain structures with the corresponding bound ligands onto their reference using TM-align^95^.

To define a final set of ligand-based druggable pockets per domain, we used a single-linkage clustering technique, merging into a single pocket all those ligands whose centroids were at a distance ≤5Å while maintaining the maximum distance between the global centroid of the cluster and the centroids of the individual compounds ≤18Å. We considered the final global cluster centroids as the pocket centroids. Overall, we found 1,604 ligand-defined pockets in 1,279 protein domains (363 unique Pfam domains in 1,205 proteins). We named this set of pockets PDB-LIG.

We then superimposed the reference PDB structure of the previous domains to their AF2 predicted counterparts by means of TM-align^95^, and transferred the location of the identified PDB-LIG pockets. We only considered those pockets having a pLDDT value >70 for all the residues at a distance ≤8Å from the pocket centroid. Overall, we identified 1,405 pockets in 1,131 domains (339 unique Pfam domains in 1,074 proteins), and named this set of pockets AF2-LIG.

#### Pocket-detection

As a complementary strategy, and to increase the overall coverage of human druggable pockets, we attempted a *de novo* identification of pockets. To establish a standardized protocol to predict them, we assessed the accuracy of different methods when identifying the PDB-LIG pockets defined above. In brief, we first removed bound ligands from reference *holo* structures (1,279 PDB structures, one per domain) and used Fpocket^70^ and P2rank^71^, two state-of-the-art methods, to detect pockets in ligand-free domain structures. Additionally, we also used Prank^71^, a functionality of P2rank aimed at rescoring the pockets predicted by Fpocket. In this way, we benchmarked three different strategies to detect and score domain binding sites. We considered that a predicted pocket and a ligand-defined pocket matched if the distance between their centroids was ≤6.14Å, which corresponded to the 95^th^ percentile of the distribution of all pairwise distances between ligand centroids within each cluster in the PDB-LIG set (**Fig S25**). We found that only 0.18% and 0.56% of Fpocket and P2rank predicted pocket pairs, respectively, had a distance between their centroids below that value. Given the apparent over-prediction of pockets of the two methods, we explored the precision/recall balance when keeping only the top scoring predicted pockets. Overall, we found that the best strategy to detect real binding sites in ligand-free structures was the combination of Fpocket detection and Prank scoring. Using the mentioned distance cut-off (6.14Å) and considering the top-2 best scored pockets for each domain, we were able to detect 72% of the real pockets while 47% of detected pockets were indeed real (**Fig 2b**).

Thus, we first ran Fpocket on the *apo* PDB reference structure for each domain to identify potential druggable pockets, we then ranked them by means of Prank, and we finally kept the top-2 ranked pockets per domain. Overall, this accounted for a total of 14,413 predicted pockets in 7,403 domains (1,806 unique Pfam domains in 4,643 proteins). We named this set of pockets PDB-PD.

We then used the same strategy and criteria as before to detect pockets onto the predicted AF2 domain structures (Fpocket and Prank combination filtering out those pockets having residues with pLDDT values <70), annotating a total of 32,202 pockets in 19,211 domains (2,409 unique Pfam domains in 10,314 proteins). We named this set of pockets AF2-PD.

For each pocket and structure, we calculated pocket volume and buriedness using rDock^62^ and BioPython^86^, respectively.

### Systematic comparison of druggable pockets within domain families

First, for each pair of pockets within the AF2-PD and AF2-LIG sets that were located at the same Pfam domains, we computed the correlation between PocketVec similarity (defined as 1-PocketVec distance), sequence identity and Cα RMSD among pockets. We selected a maximum of 10 protein domains per Pfam domain to avoid biasing the results towards the most frequently occurring ones. We then removed those domain pairs having a global TM-score <0.5 and we performed all pairwise residue mappings using the corresponding Pfam multiple sequence alignments (MSA). After that, pockets (residues <8Å) were compared on the basis of their sequences and structures (sequence identity and Cα RMSD, respectively). In fact, we only considered those pocket pairs having a sequence alignment coverage ≥80% and a centroid distance <6.14Å after domain structural alignment, to account for structural variability (see *Pocket detection* in the previous section).

Additionally, we also ran an all-against-all comparison of pockets in the human pocketome (e.g. PDB-LIG, PDB-PD, AF2-LIG and AF2-PD). Global domain structures were compared using TM-align^95^ (Cα RMSD, TM-score), while domain sequence identities were calculated using global pairwise alignments in BioPython^86^ (Needleman-Wunsch algorithm, BLOSUM62, gap opening = −10, gap extension = −0.5).

### Comparison of kinase inhibition profiles with PocketVec descriptors, sequence and structure similarity measurements

We collected experimentally determined binding affinities from Klaeger et al.^77^ and Reinecke et al.^78^ (**Table S2**). We considered undefined measures as inactive, and low-confidence and high-confidence measures were binarized at 30 nM (as recommended in Pharos^96^). We then removed all compounds that did not inhibit any kinase and all protein kinases that did not have any inhibitor or any PocketVec descriptor in the Protein Kinase Domain (Pfam PF00069). In this way, we eventually defined a binary inhibition matrix between 111 protein kinases and 94 small molecule kinase inhibitors for Klaeger et al.^77^ (**Fig 5a** and **Table S2a**) and between 73 kinases and 164 inhibitors for Reinecke et al.^78^ (**Fig 5b** and **Table S2b**).

We pairwise compared protein kinases on the basis of their binarized inhibition profiles (Jaccard similarity) and their PocketVec descriptors (employing the minimum distance among all PocketVec descriptors within their Protein Kinase Domains PF00069). Additionally, we also performed sequential and structural comparisons between kinases at domain level following the same strategy as in previous sections. In brief, domain structures were compared using TM-align^95^ (Cα RMSD, TM-score) and domain sequence identities were calculated using global pairwise alignments in BioPython^86^ (Needleman-Wunsch algorithm, BLOSUM62, gap opening =−10, gap extension = −0.5). Only the highest TM-score and sequence identity values among domains were considered for each pair of kinases.

## Acknowledgements

P.A. acknowledges the support of the Generalitat de Catalunya (RIS3CAT Emergents CECH: 001-P-001682 and VEIS: 001-P-001647; and 2021 SGR 00876), the Spanish Ministerio de Ciencia, Innovación y Universidades (PID2020-119535RB-I00), the Instituto de Salud Carlos III (IMPaCT-Data), and the European Commission (RiPCoN: 101003633). A.C-C. is a recipient of an FI fellowship (2020 FI_B 00094). We also acknowledge institutional funding from the Spanish Ministry of Science and Innovation through the Centres of Excellence Severo Ochoa Award, and from the CERCA Programme / Generalitat de Catalunya.

