## Supplementary Information for "Comprehensive detection and characterization of human druggable pockets through novel binding site descriptors"

### Supplementary Material

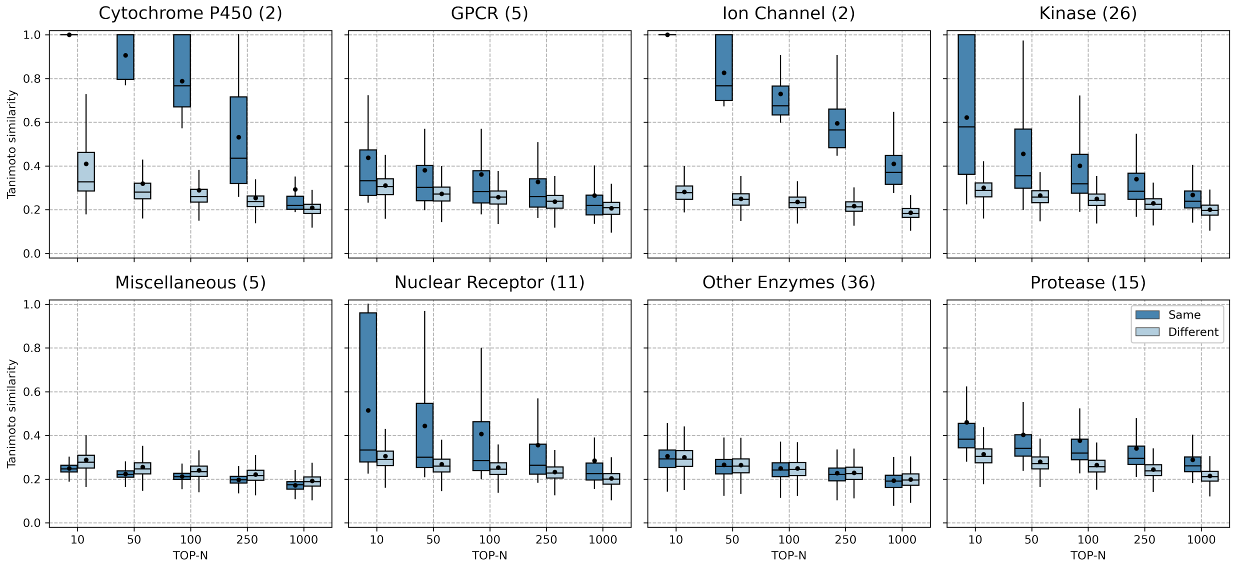


**Fig S1**: Similar proteins bind similar active compounds. For each protein family (e.g. Cytochrome P450, GPCR, etc.) we compared all active compounds between protein pairs from the same family (Same) and from different families (Different). Compounds were compared on the basis of their Tanimoto Similarities (y-axis, ECFP4 with 2048 bits using RDKit – https://rdkit.org) and only the TOP-N (x-axis: 10, 50, 100, 250, 1000) highest similarities were considered per each family. All distributions of Same and Different categories are significantly different (Mann-Whitney U test, pvalue <0.05, alternative *greater* using SciPy[1]) except Miscellaneous (10, 50, 100, 250, 1000) and Other Enzymes (10, 50, 100, 250, 1000). Box plots indicate median (middle line), 25th, 75th percentile (box), and max and min value within the 1.5*25th and 1.5*75th percentile range (whiskers). Black dots represent mean values. The number of proteins within each family is specified in the title in parenthesis. Data was downloaded from http://dude.docking.org in January 2023.

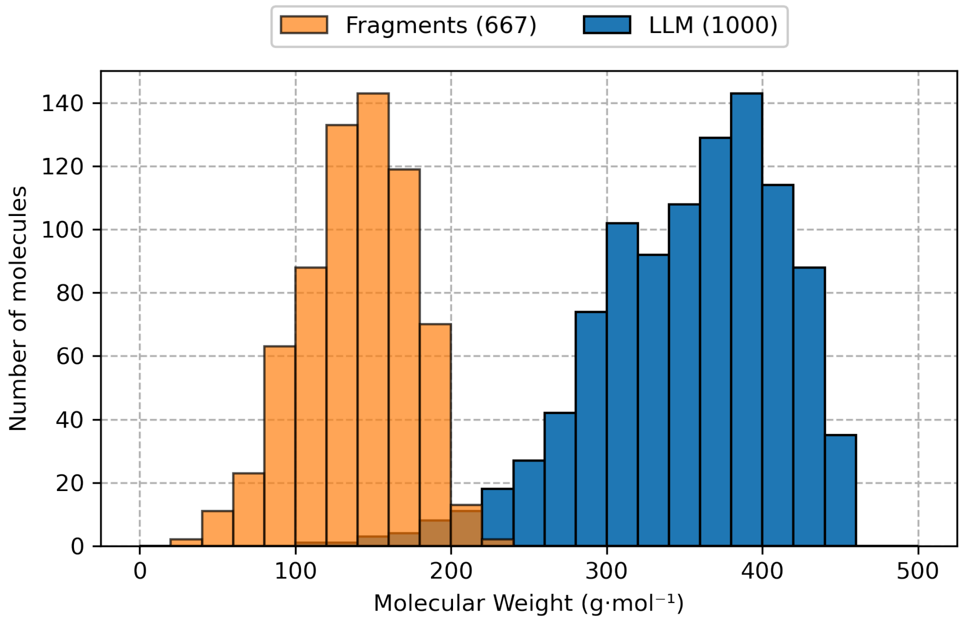


**Fig S2**: Histograms of molecular weights (x-axis) from fragments (667, orange) and lead-like molecules (1,000, blue).

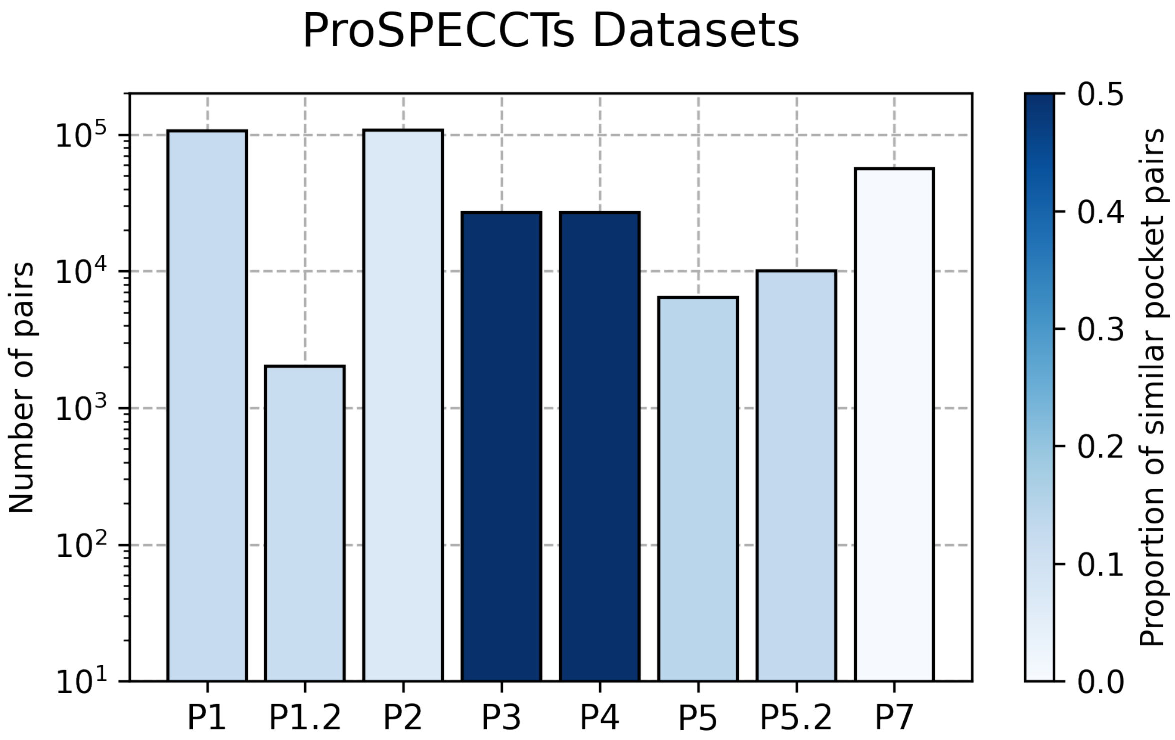


**Fig S3**: ProSPECCTs overview. Each bar represents a ProSPECCT dataset (x-axis) and indicates the number of protein-ligand binding site pairs (y-axis) together with the proportion of similar pocket pairs within each dataset (color tone). Please, see the *Online Methods* section for a detailed description of each dataset.

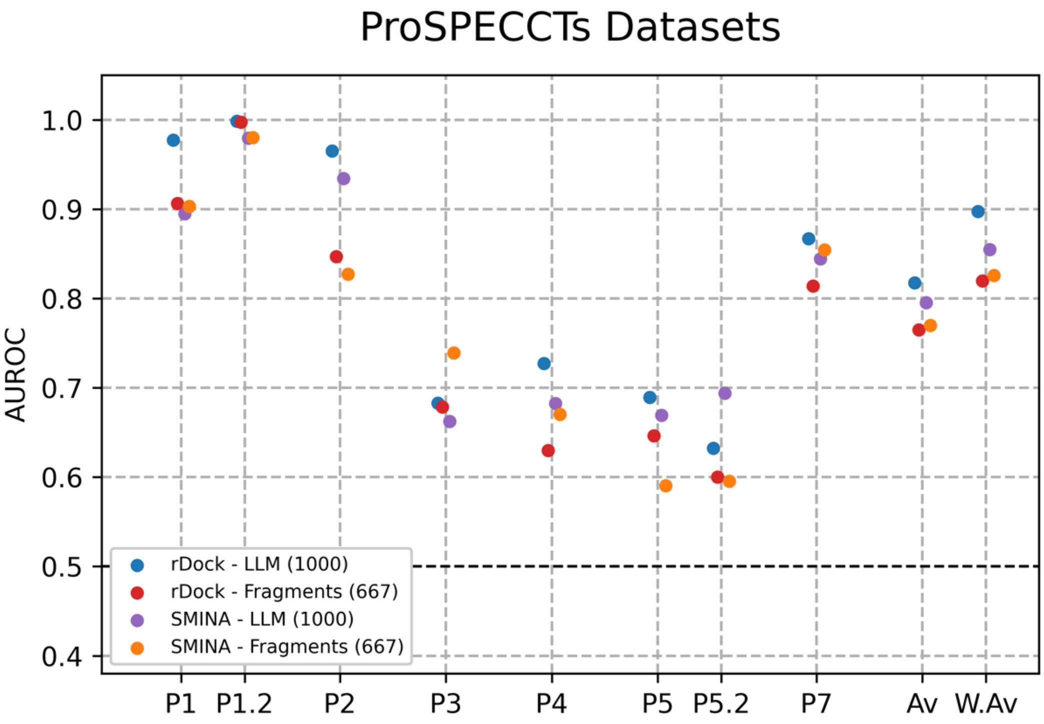


**Fig S4**: Performances (AUROC, y-axis) among ProSPECCTs datasets (x-axis). Each colored dot represents the individual performance of our strategies with all possible combinations of docking approaches (rDock and SMINA) and compound collections (1,000 LLM and 667 fragments). Av. values represent the average performance among ProSPECCTs datasets for each individual strategy and W. Av. values weight the average value according to the number of pairs within each dataset.

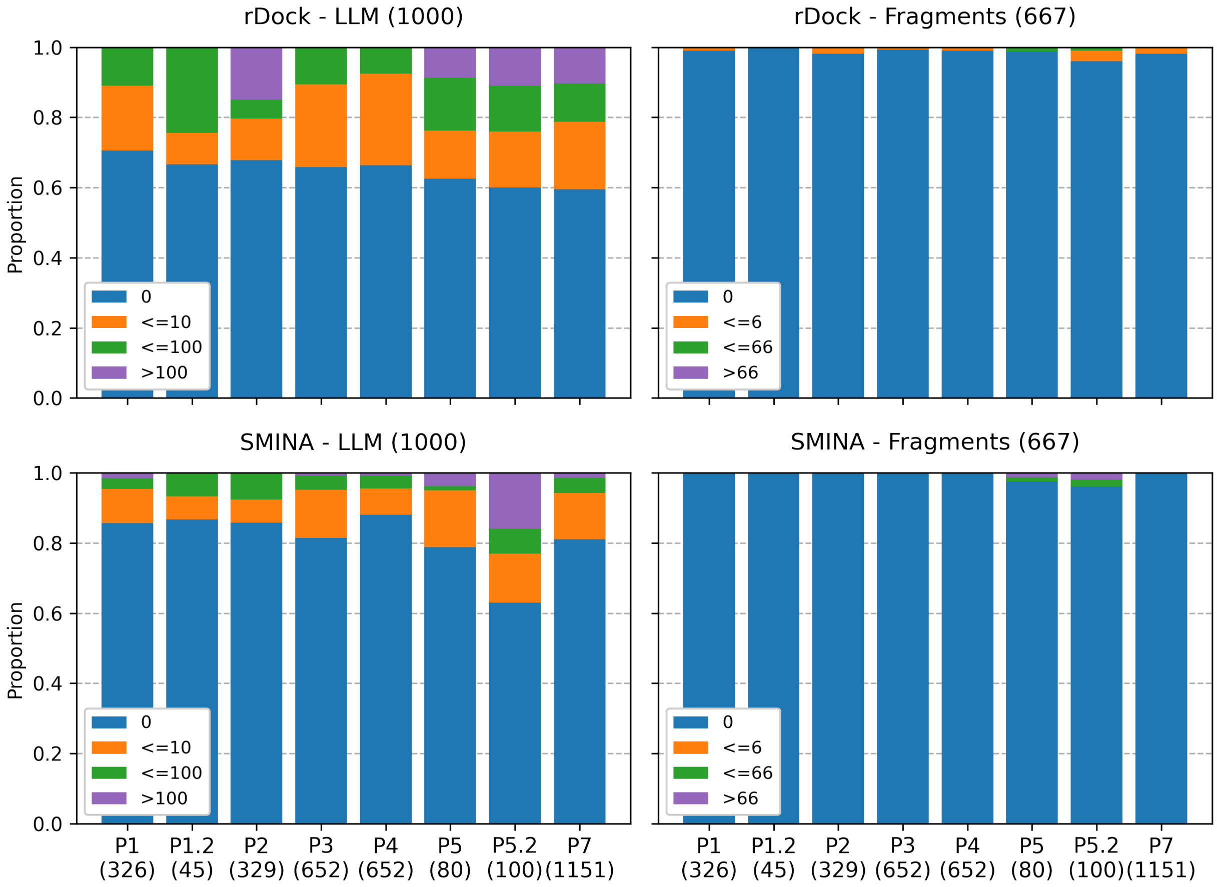


**Fig S5**: Proportion of outlier molecules. Colored bars represent the proportion of outlier molecules (y-axis) within each ProSPECCTs dataset (x-axis) for each combination of docking method (rDock and SMINA) and compound collection (1,000 LLM and 667 fragments). Colors indicate the number of outlier molecules assigned to each structure. The number of structures per dataset is specified in parenthesis (x labels).

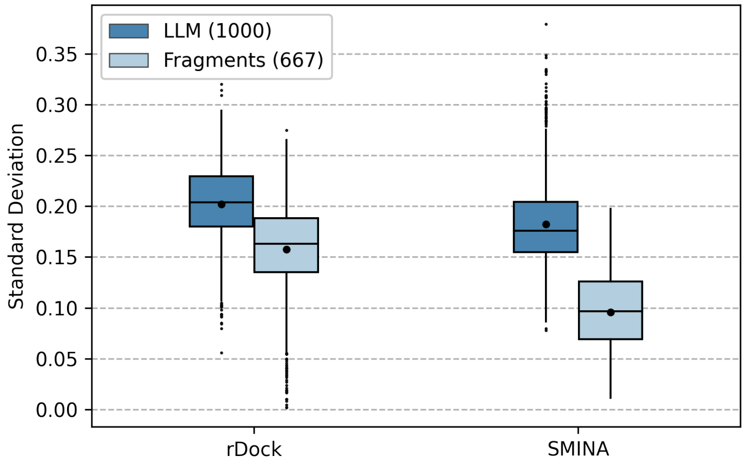


**Fig S6**: Ranking diversity differences. For each docking method (rDock and SMINA) and compound collection (1,000 LLM and 667 fragments) ProSPECCTs pockets were considered all at once (3,335 pockets) and the standard deviation (in terms of rankings) was calculated for each tested compound. For both docking strategies standard deviation distributions between LLM and fragments are statistically different (Mann-Whitney U test, $p<10^{-70}$ in both cases). Box plots indicate median (middle line), 25th, 75th percentile (box), and max and min value within the 1.5*25th and 1.5*75th percentile range (whiskers). Black dots represent mean values.

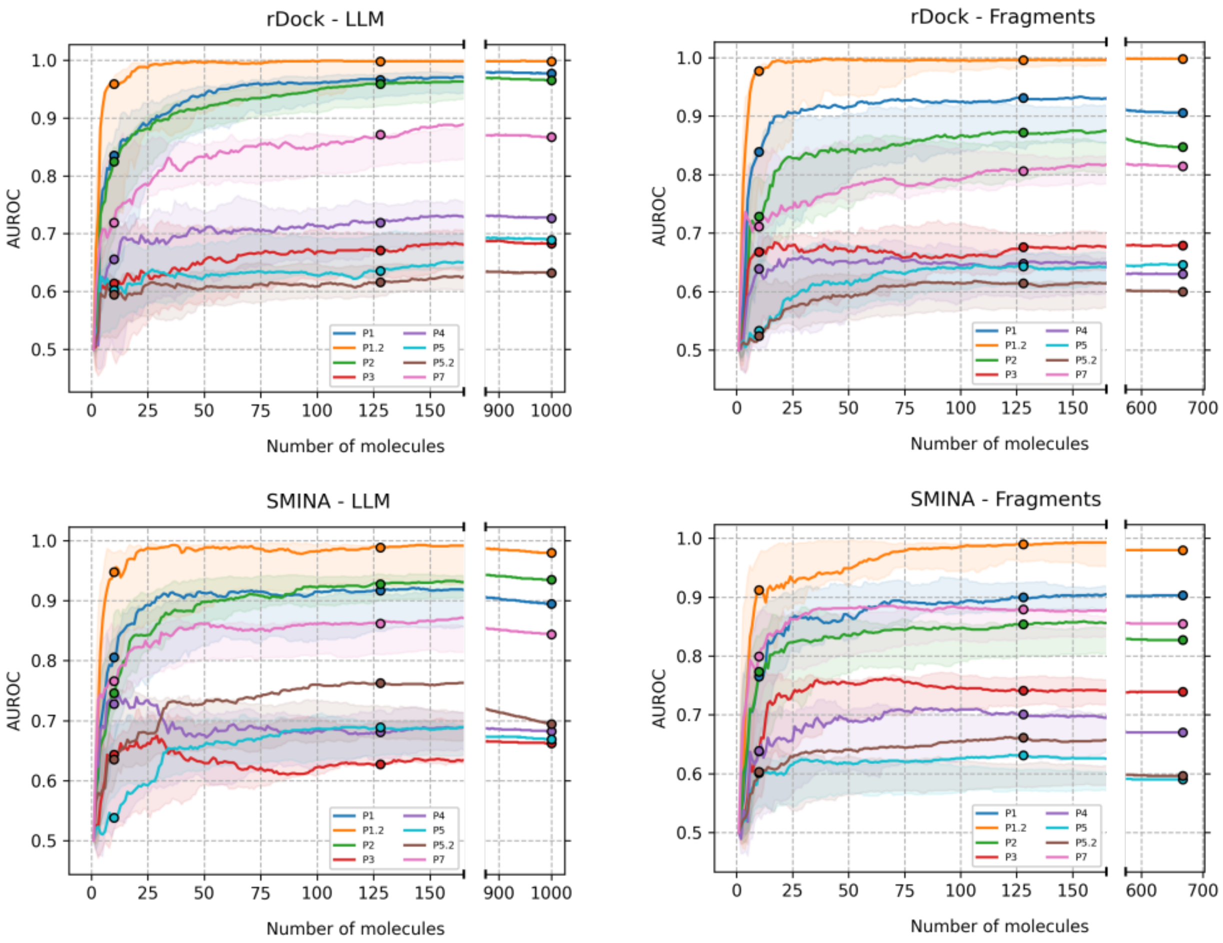


**Fig S7**: Performances (AUROC, y-axis) of our descriptors among ProSPECCTs datasets (color, P6 and P6.2 not included. For further details, please see *Online Methods*) generated with a varying number of predefined compounds (x-axis, from 1 to complete sets, sorted by entropy). Colored points indicate the performance at 10, 128 and 1,000/667 molecules. Colored areas represent the best and worst performances obtained with randomly (x100) selected compounds (not sorted by entropy values). Each subplot corresponds to a combination of docking method (rDock and SMINA) and compound collection (1,000 LLM and 667 fragments).

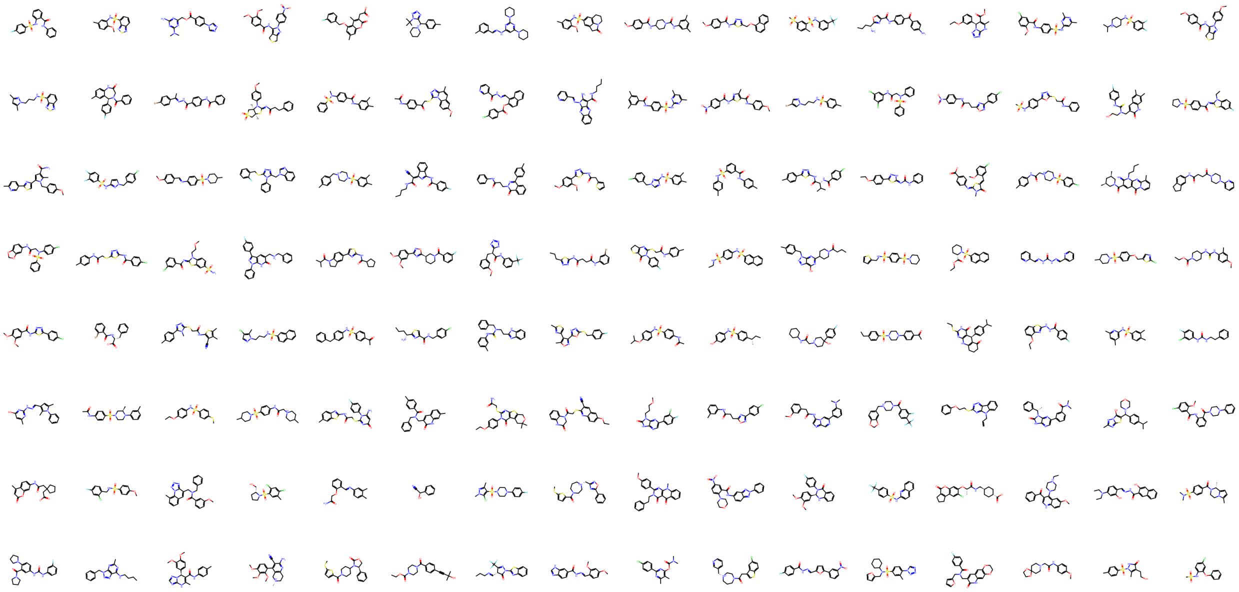

**Fig S8**: Standard 128 lead-like molecules used to generate PocketVec descriptors.

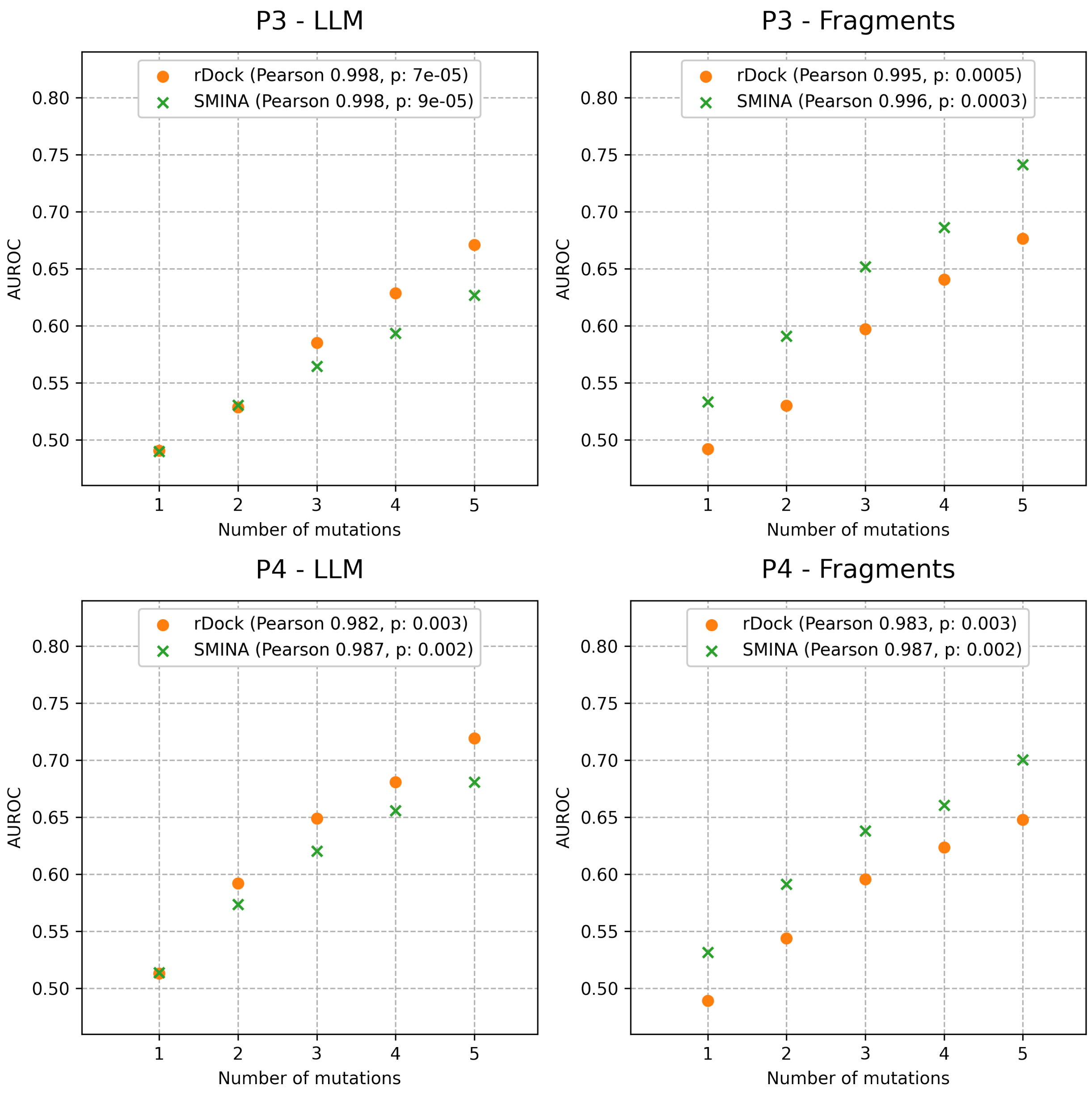


**Fig S9**: Correlation between the AUROC (y-axis) and the number of artificial mutations (x-axis) in pocket-lining residues leading to changes in physicochemical properties (ProSPECCTs P3) and both physicochemical and shape properties (ProSPECCTs P4). Each subplot corresponds to a specific ProSPECCTs dataset (P3, P4) and collection of compounds (128 LLM and 128 fragments). Dot color and marker indicate the used docking method (rDock for rigid docking and SMINA for flexible docking). In the legend, the corresponding Pearson's Correlation Coefficients are reported.

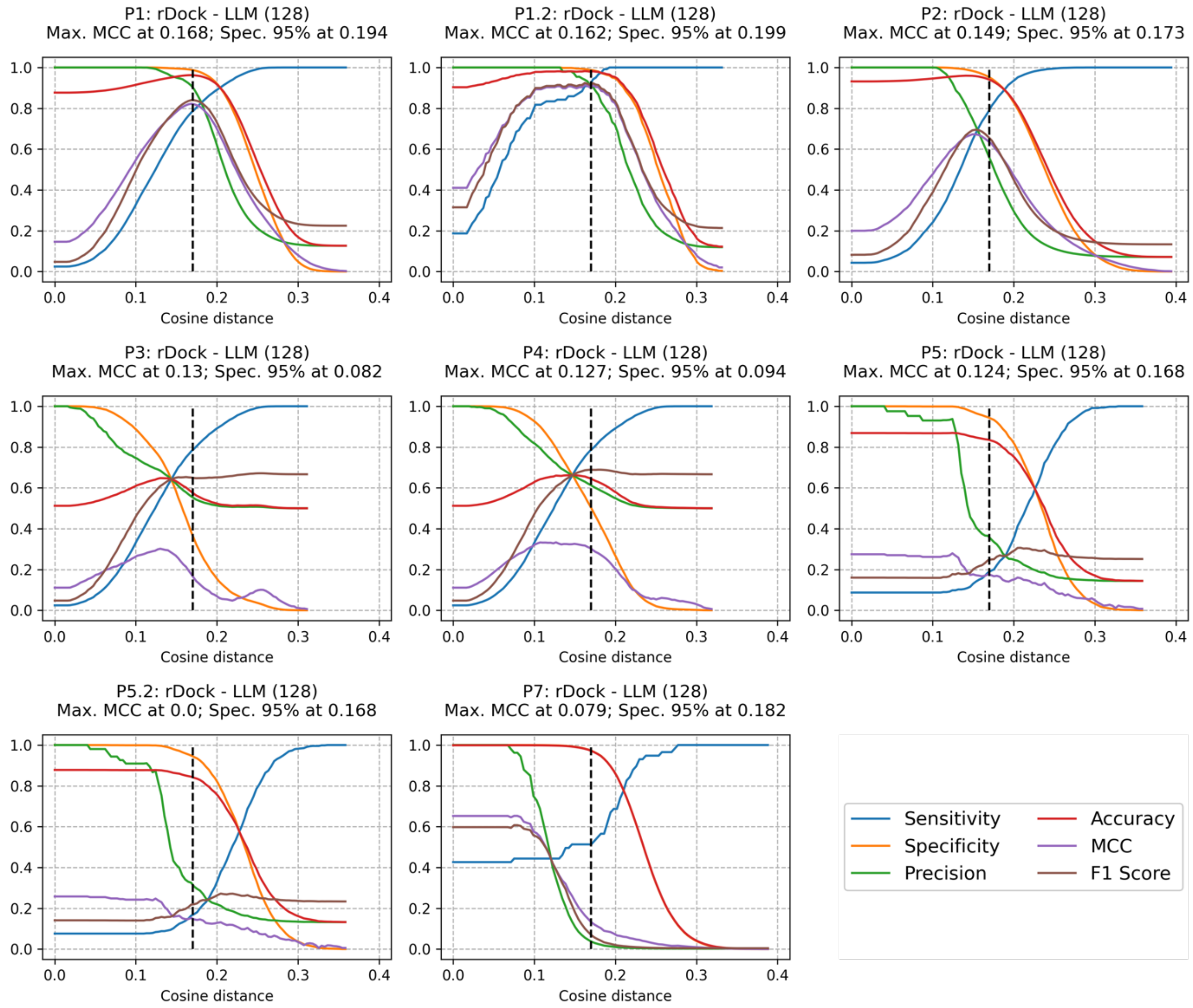


**Fig S10**: Evolution of (y-axis) Sensitivity (blue), Specificity (orange), Precision (green), Accuracy (red), Matthew's Correlation Coefficient (MCC, purple) and F1 Score (brown) at multiple values of PocketVec cosine distance (x-axis, rDock as docking method and 128 LLM as collection of compounds). Each subplot corresponds to a ProSPECCTs dataset (see title).

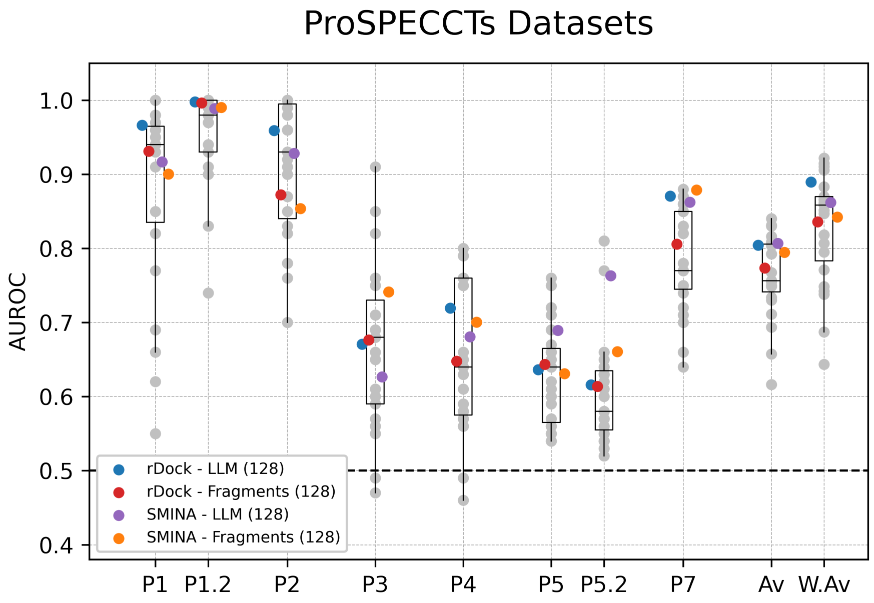


**Fig S11**: Performances (AUROC, y-axis) of distinct approaches to compare binding sites among ProSPECCTs datasets (x-axis). Gray dots represent the individual performances of existing strategies to compare pockets (not only those based on pocket descriptors). Box plots indicate median (middle line), 25th, 75th percentile (box), and max and min value within the 1.5*25th and 1.5*75th percentile range (whiskers). Colored dots represent the individual performances of our strategies with all possible combinations of docking approaches (rDock and SMINA) and reduced sets of compound collections (128 LLM and 128 fragments). Av. values represent the average performance among ProSPECCTs datasets for each individual method and W. Av. values weight the average value according to the number of pairs within each dataset.

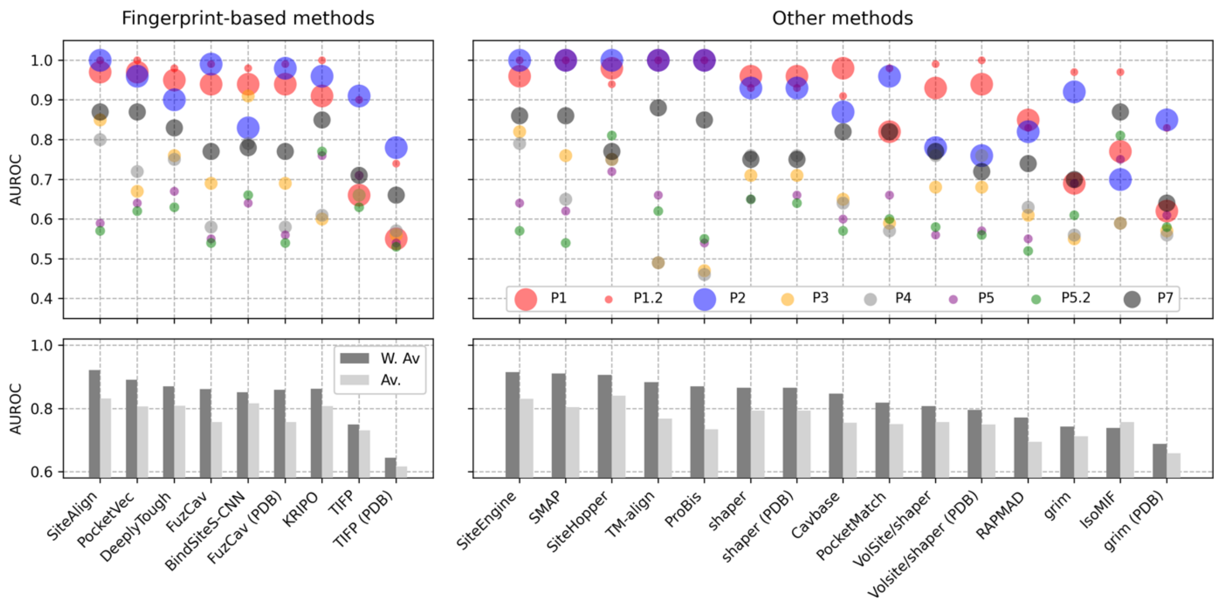


**Fig S12**: Exhaustive comparison of different methods used to assess pocket similarity among ProSPECCTs datasets. Left plots represent the performances of strategies based on the generation of pocket fingerprints or descriptors. The superior plot indicates the individual performance (y-axis, AUROC) of each method (x-axis) within each dataset (dot color, dot size is proportional to dataset size -number of pocket pairs). The plot below indicates the average and weighted average performance (according to the number of pairs within each dataset) of each method. Methods are sorted by W.Av values. Right plots represent the same information as mentioned before but for those existing strategies not based on pocket fingerprinting.

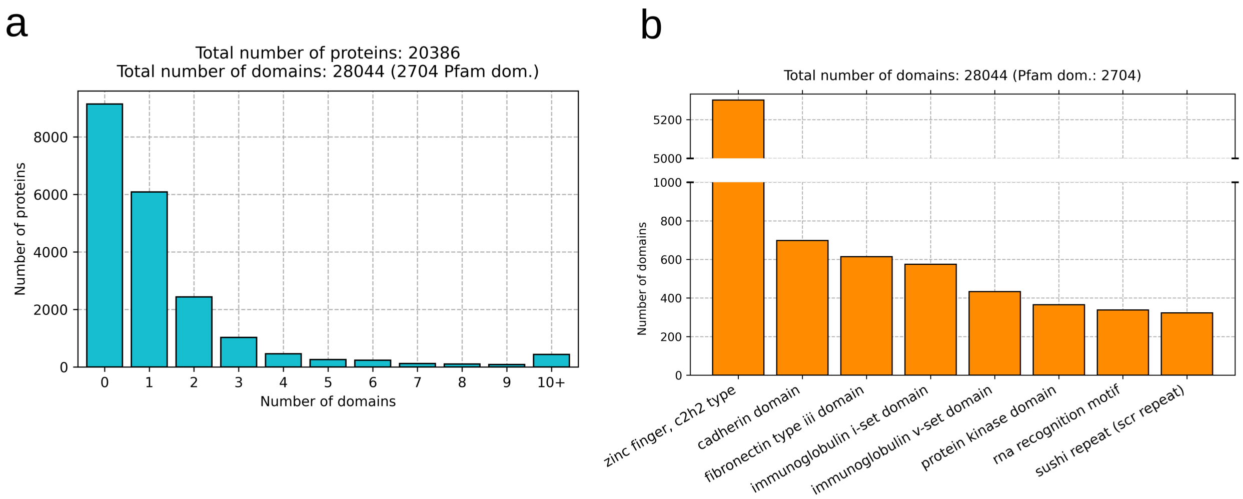


**Fig S13**: **a)** Number of domains per protein. Each bar corresponds to the number of human proteins (y-axis) having the specified number of domains (x-axis). **b)** Number of domain occurrences (y-axis) per Pfam domain (x-axis).

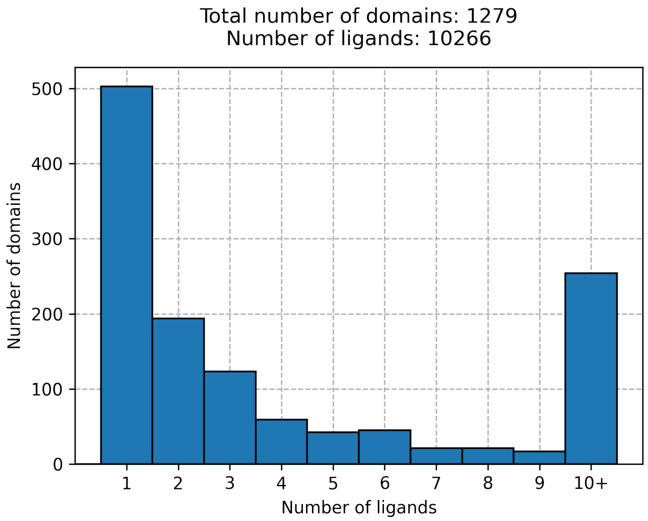


**Fig S14**: Number of ligands per domain. Each bar corresponds to the number of human protein domains (y-axis) having the specified number of ligands (x-axis) fulfilling our criteria (see *Online Methods*).

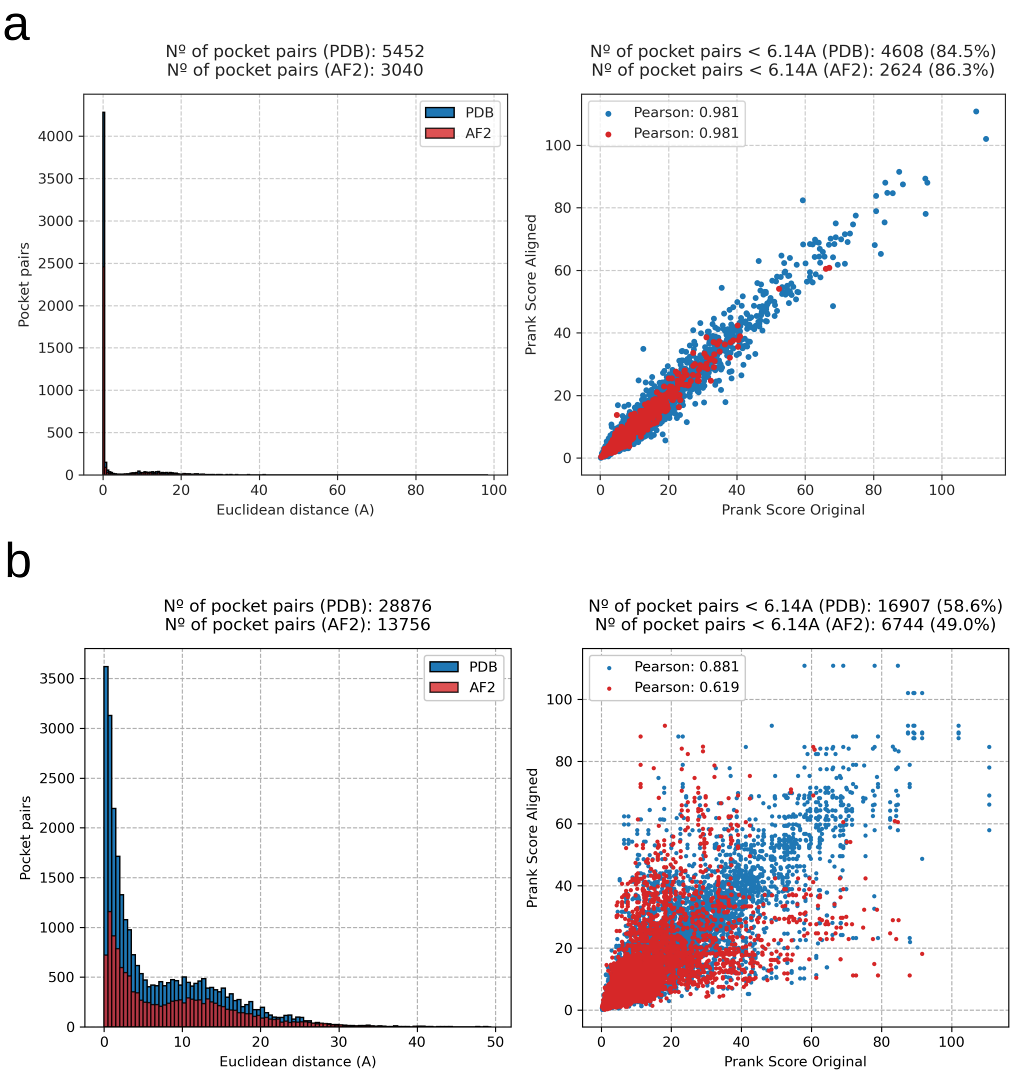


**Fig S15**: Evaluating pocket detection consistency in PDB and AF2 structures. First, we randomly selected 1,000 domains from the PDB-PD set. We only considered those domains having more than one PDB structure (786) and selected a single structure as the reference one. Then, the remaining PDB structure(s) and the corresponding AF2 model were superposed against the reference structure using TM-align[2]. **a)** Pockets (top-2) were detected in original and aligned PDB and AF2 structures. Original pocket centroids were superposed against aligned pocket centroids using the corresponding translation and rotation matrices. Minimum Euclidean distances between aligned and original pockets were calculated and are shown in a histogram representation (left), distinguishing between PDB and AF2 structures. For those pocket pairs that were consistently detected (distance ≤6.14Å between aligned and original structures, see *Online Methods*), the correlation between their original and aligned scoring is shown on the right plot, together with the corresponding Pearson's correlation coefficients. **b)** Pockets (top-2) were detected in aligned PDB and AF2 structures (including the reference one). Minimum Euclidean distances between aligned (PDB and AF2) and reference pockets were calculated and are shown in a histogram representation (left), distinguishing between PDB and AF2 structures. For those pocket pairs that were consistently detected (distance ≤6.14Å between aligned and original structures), the correlation between their aligned (PDB and AF2) and reference scoring is shown on the right plot, together with the corresponding Pearson's Correlation Coefficients.

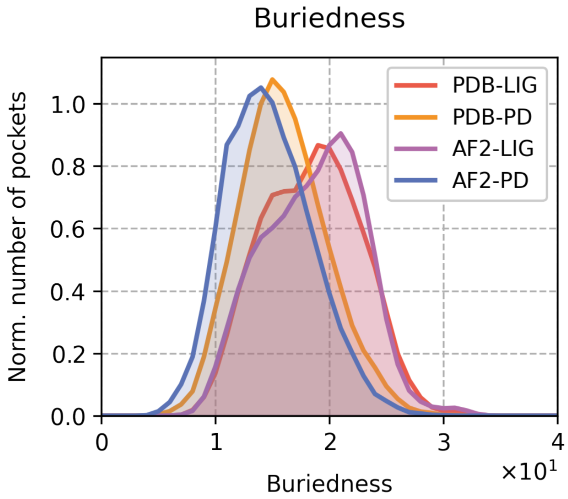


**Fig S16**: Distribution of buriedness values (x-axis, buriedness is defined as the number of protein residues having a distance below 8Å to the pocket centroid) for each pocket set: PDB-LIG (1,604 pockets, red), PDB-PD (14,413 pockets, orange), AF2-LIG (1,405 pockets, purple) and AF2-PD (32,302 pockets, blue). All distributions are normalized (density, y-axis). The average buriedness values are 18.5, 16.0, 18.6 and 14.7, respectively.

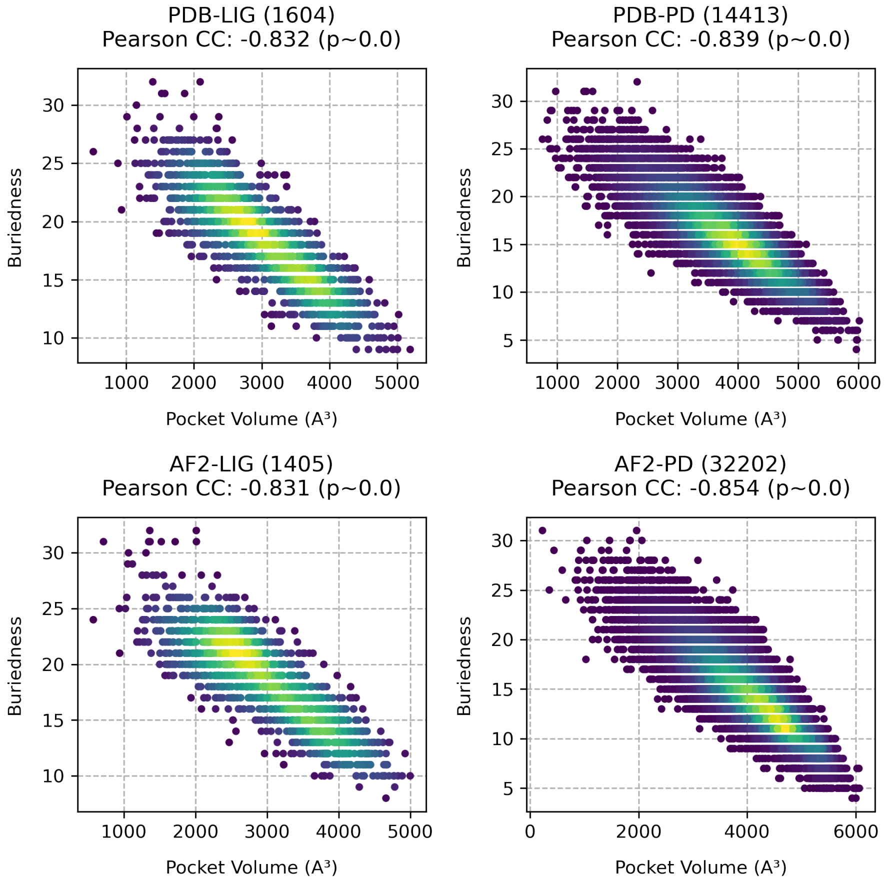


**Fig S17**: Correlation between pocket volume and buriedness (buriedness is defined as the number of protein residues having a distance below 8Å to the pocket centroid). For each pocket set (PDB-LIG, PDB-PD, AF2-LIG and AF2-PD) every point represents a pocket in terms of its volume (x-axis) and buriedness value (y-axis). Points are colored by density, and Pearson's Correlation Coefficients are reported in the title.

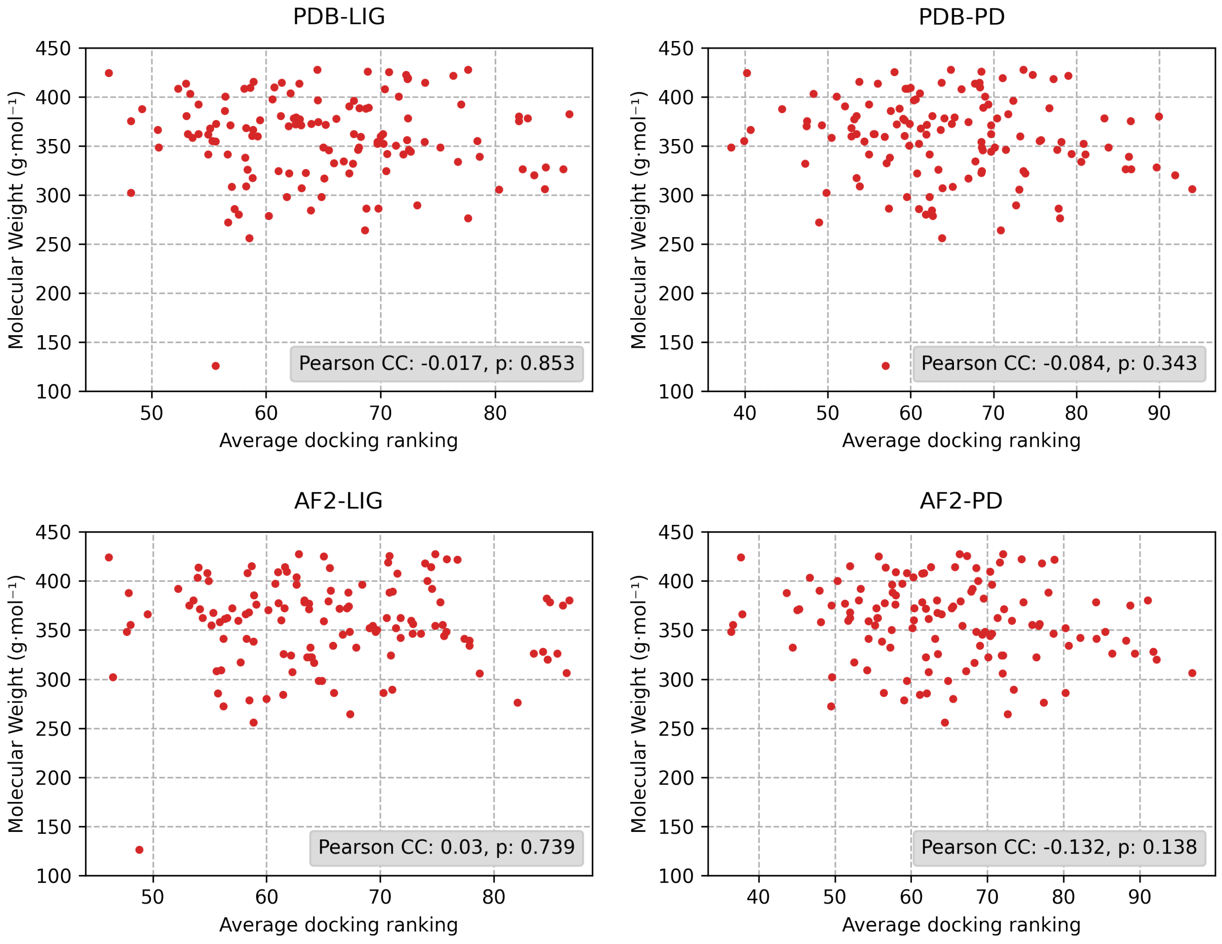


**Fig S18**: Correlation between docking ranking and molecular weight. For each pocket set (PDB-LIG, PDB-PD, AF2-LIG and AF2-PD), each point represents a lead-like molecule in terms of its average docking ranking (x-axis) within the set and its molecular weight (y-axis). Pearson's Correlation Coefficients (and p-values) are included in the legend.

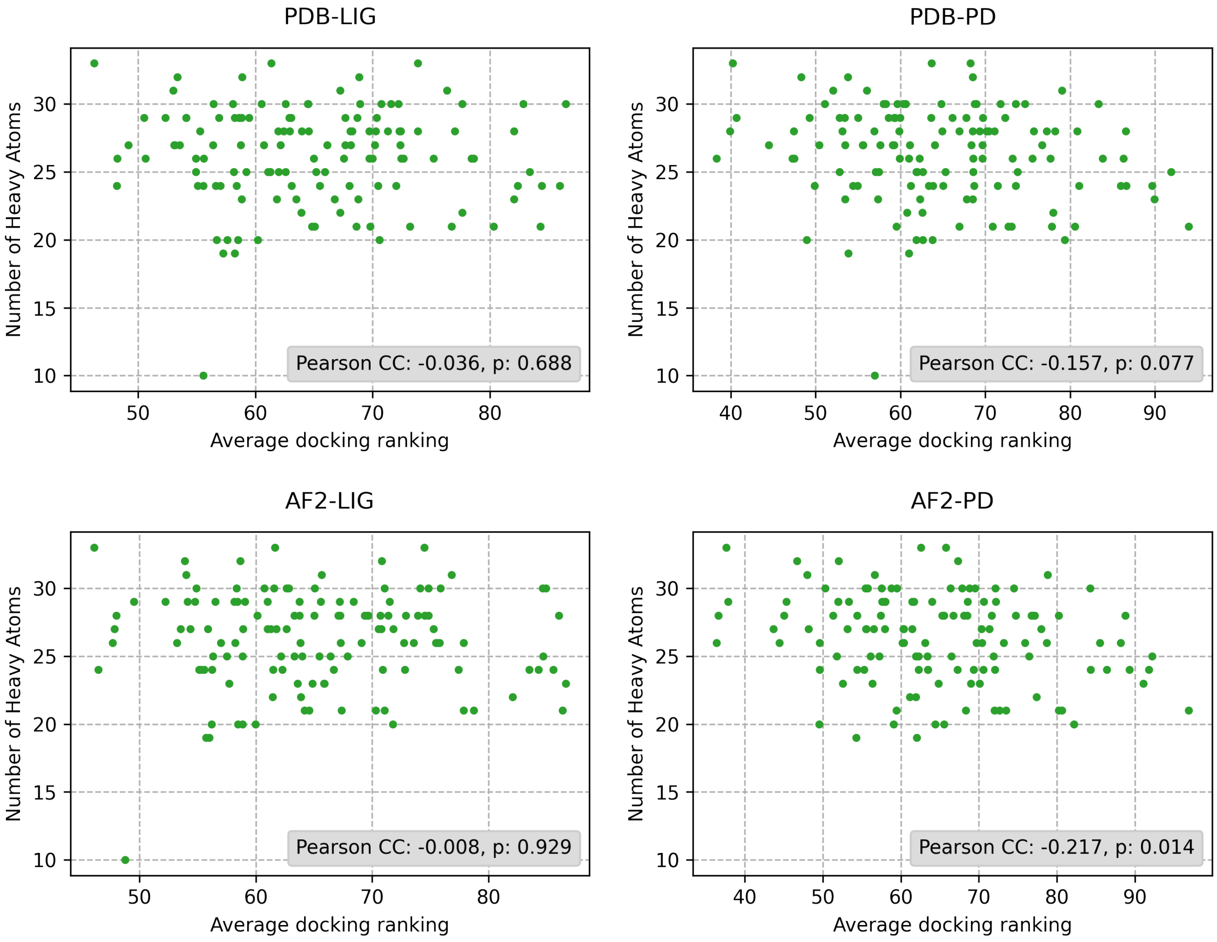


**Fig S19**: Correlation between docking ranking and number of heavy atoms. For each pocket set (PDB-LIG, PDB-PD, AF2-LIG and AF2-PD), each point represents a lead-like molecule in terms of its average docking ranking within the set (x-axis) and its number of heavy atoms (y-axis). Pearson's Correlation Coefficients are included in the legend.

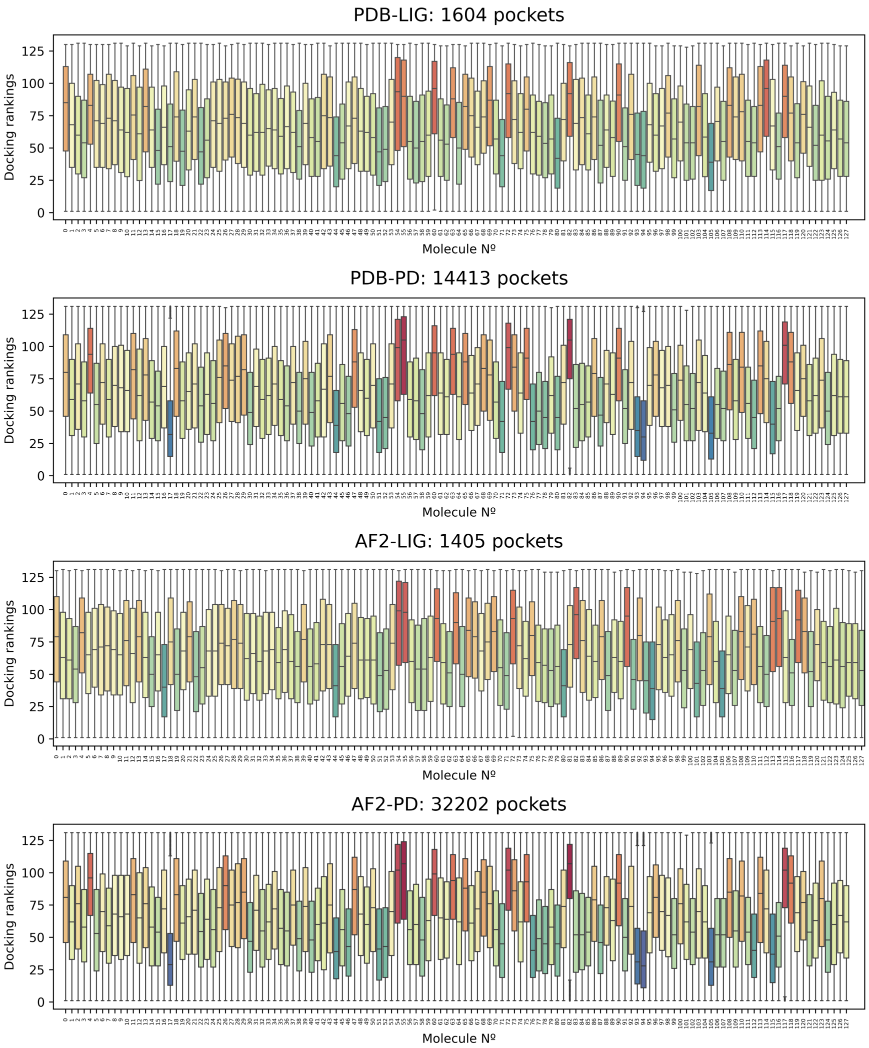


**Fig S20**: For each pocket set (PDB-LIG, PDB-PD, AF2-LIG and AF2-PD), docking rankings for each lead-like molecule (x-axis. Molecules are sorted in a predefined order) are represented as a box plot (y-axis). Box plots indicate median (middle line), 25th, 75th percentile (box), and max and min value within the 1.5*25th and 1.5*75th percentile range (whiskers) and are colored according to median ranking values, from higher (red) to lower (blue) values.

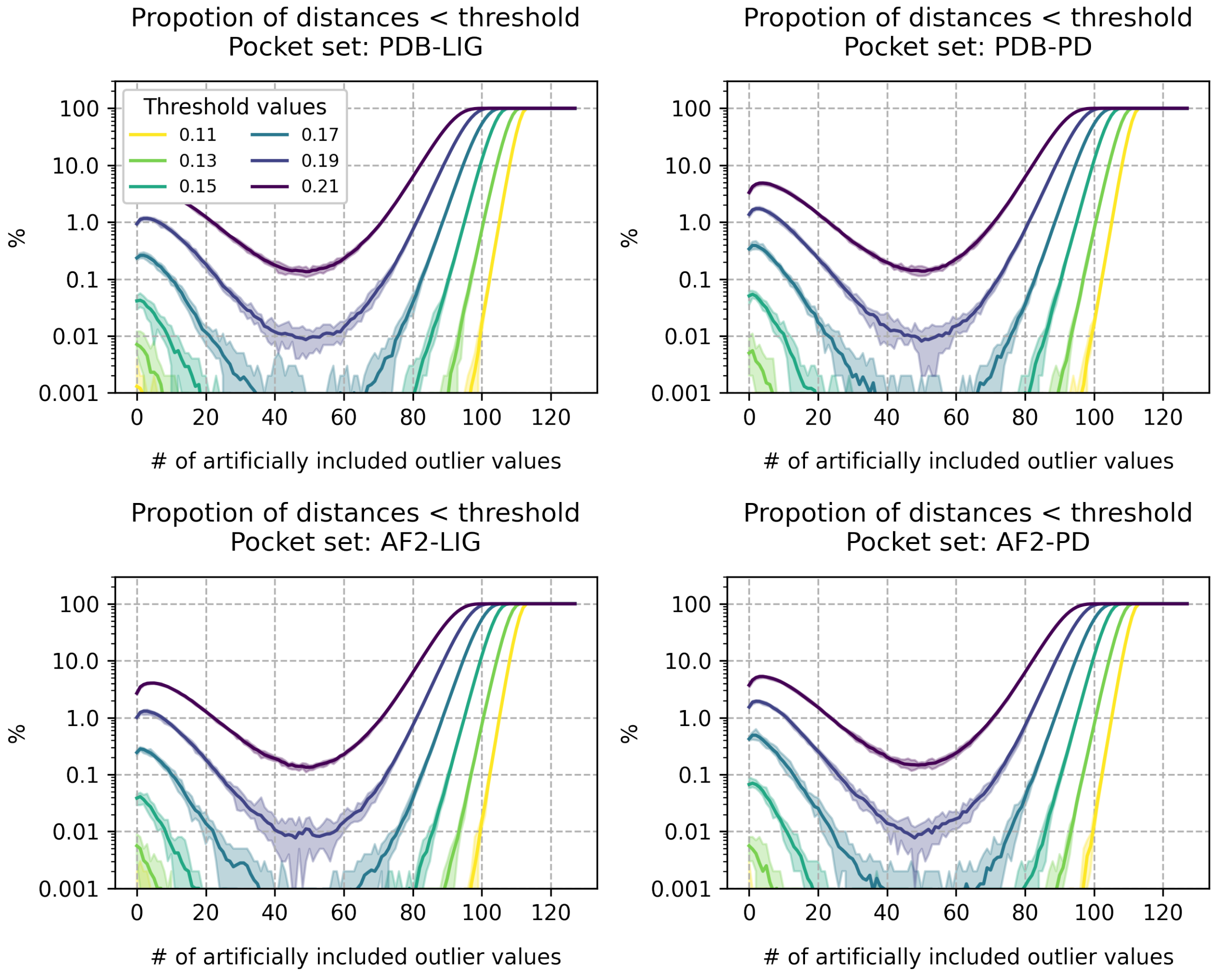


**Fig S21**: For each pocket set (PDB-LIG, PDB-PD, AF2-LIG and AF2-PD), proportion of originally dissimilar pocket pairs (y-axis, distances >threshold) classified as similar (distances <threshold) due to the artificial insertion of a growing number of outlier values (x-axis) at different distance thresholds. To run the analysis, 100,000 PocketVec distances were randomly sampled (x10 times) from each pocket set. Solid lines represent average results and shadowed areas correspond to minimum and maximum proportions among samples.

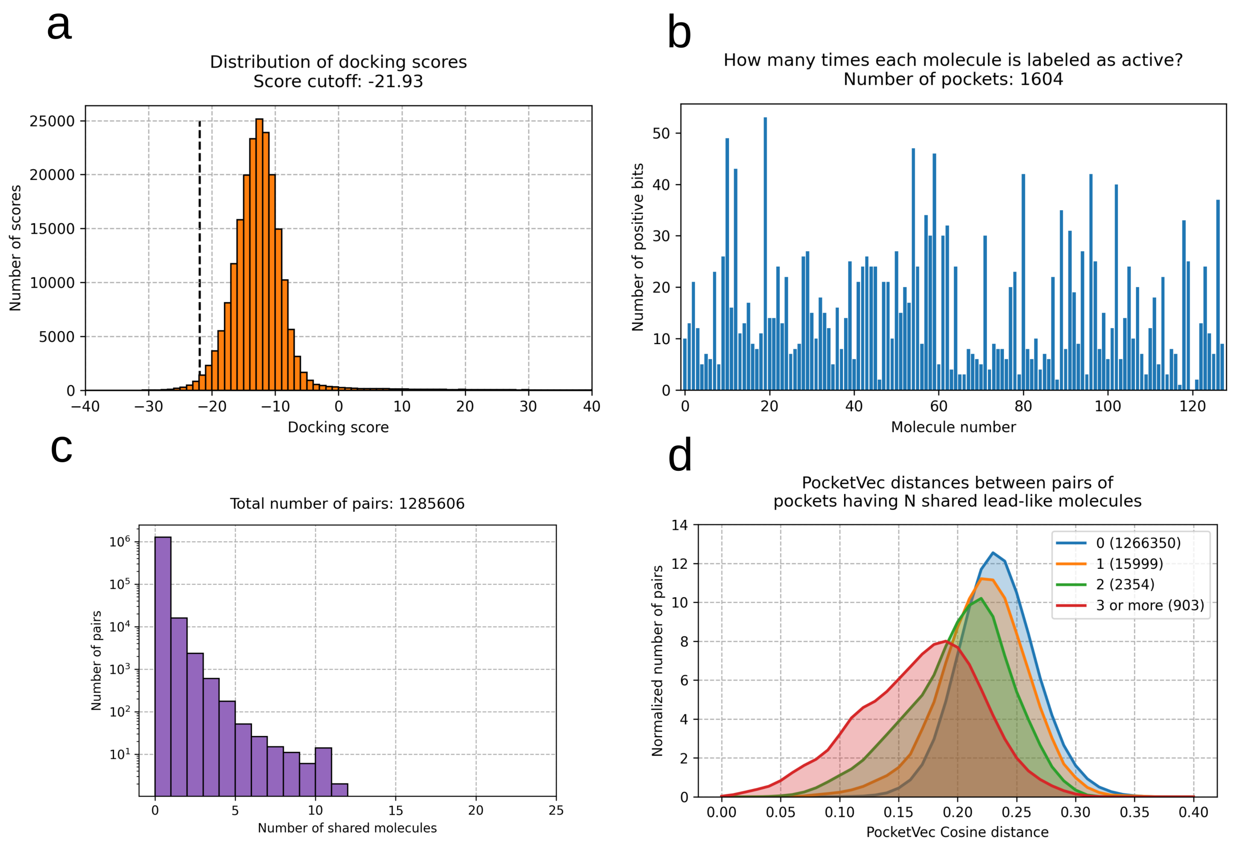


**Fig S22**: Computational evaluation of the hypothesis that pockets sharing ligands have lower PocketVec distances. **a)** Distribution (y-axis) of rDock docking scores (x-axis) from the standard set of 128 lead-like molecules (see *Methodological development and implementation*) against 1,604 PDB-LIG pockets. The highlighted cut-off score (-21.94) corresponds to the 1$^{st}$ percentile of the distribution and was used to label all pocket-compound pairs as active (<-21.94) or inactive (>-21.94). **b)** Number of PDB-LIG pockets (y-axis) for which each lead-like molecule (x-axis, from molecule #1 to molecule #128 sorted in a predefined order) was labeled as active. **c)** Number of pocket pairs (y-axis) per number of shared lead-like molecules (x-axis). **d)** Distributions (density, y-axis) of PocketVec cosine distances (x-axis) grouped by the number of shared lead-like molecules between pockets (0, 1, 2 and 3 or more). The number of pocket pairs per number of shared lead-like molecules is specified in parenthesis.

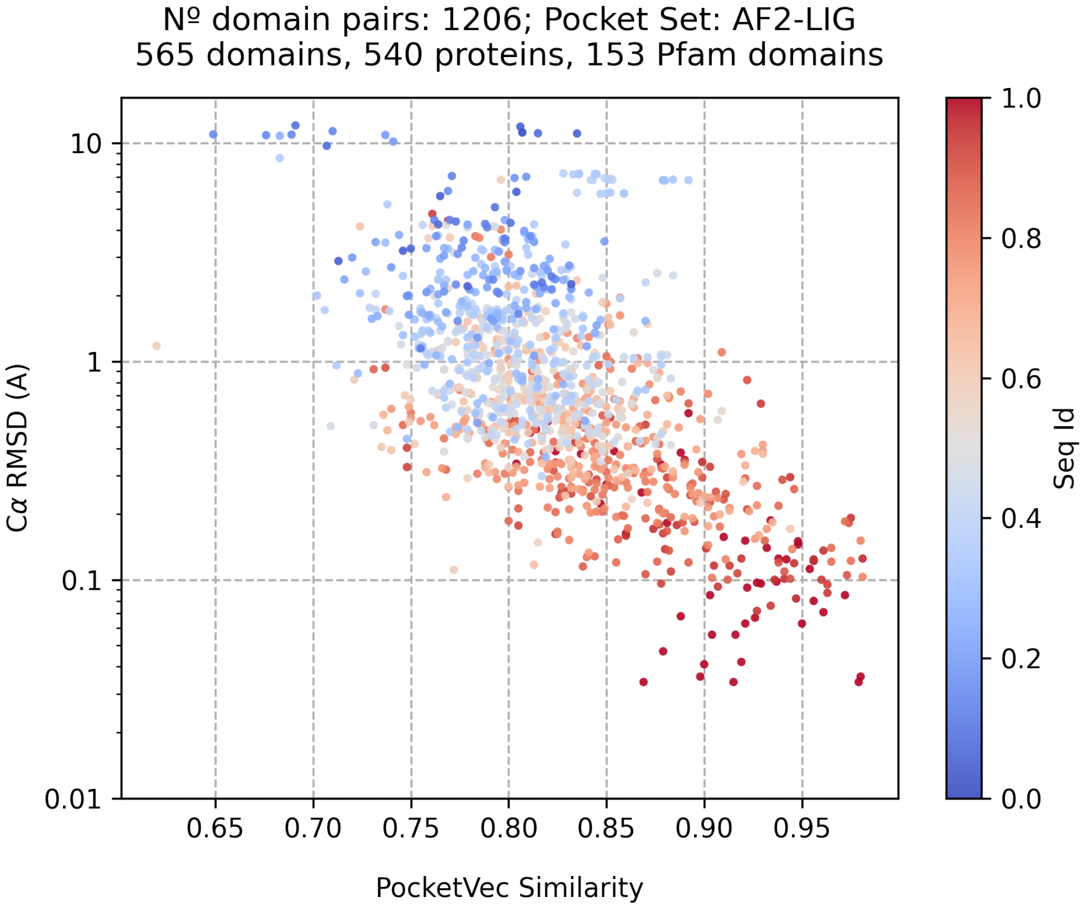


**Fig S23**: Correlation between PocketVec similarity (x-axis, defined as 1-PocketVec distance), structural similarity (y-axis, Cα RMSD) and sequence identity (color) among pockets located at the same Pfam domain (max. 10) in the AF2-LIG pocket set. Pearson CC between PocketVec similarity and sequence identity: 0.57 (p-value $<10^{-100}$). Pearson CC between PocketVec similarity and RMSD: -0.35 (p-value $<10^{-35}$).

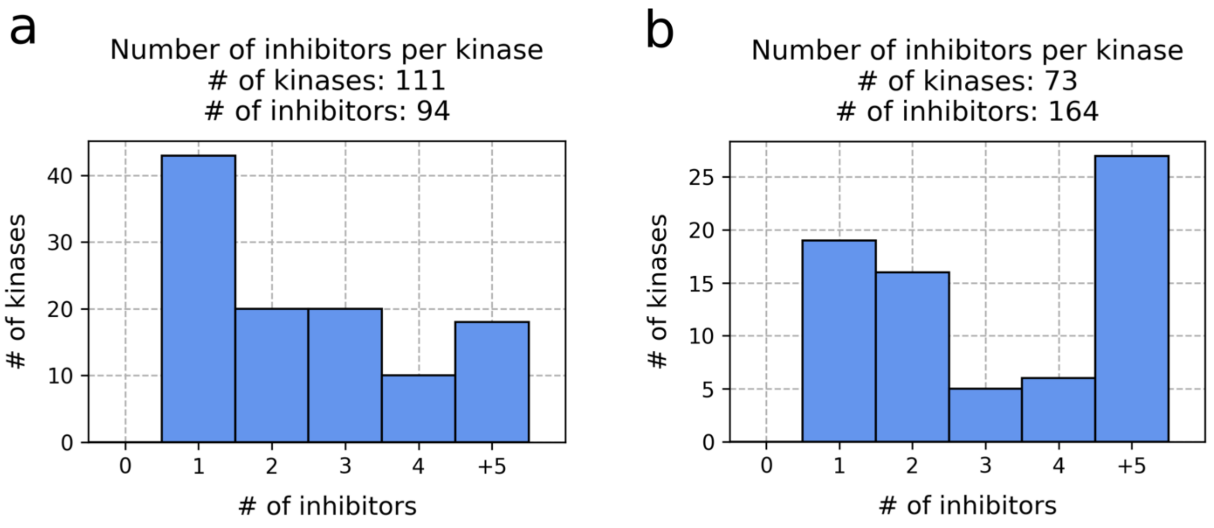


**Fig S24**: Number of inhibitors per kinase obtained from Kleager et al. (2017 set)[3] and Reinecke et al. (2023 set)[4] binarized at 30 nM. Each bar corresponds to the number of kinases (y-axis) having the specified number of inhibitors (x-axis).

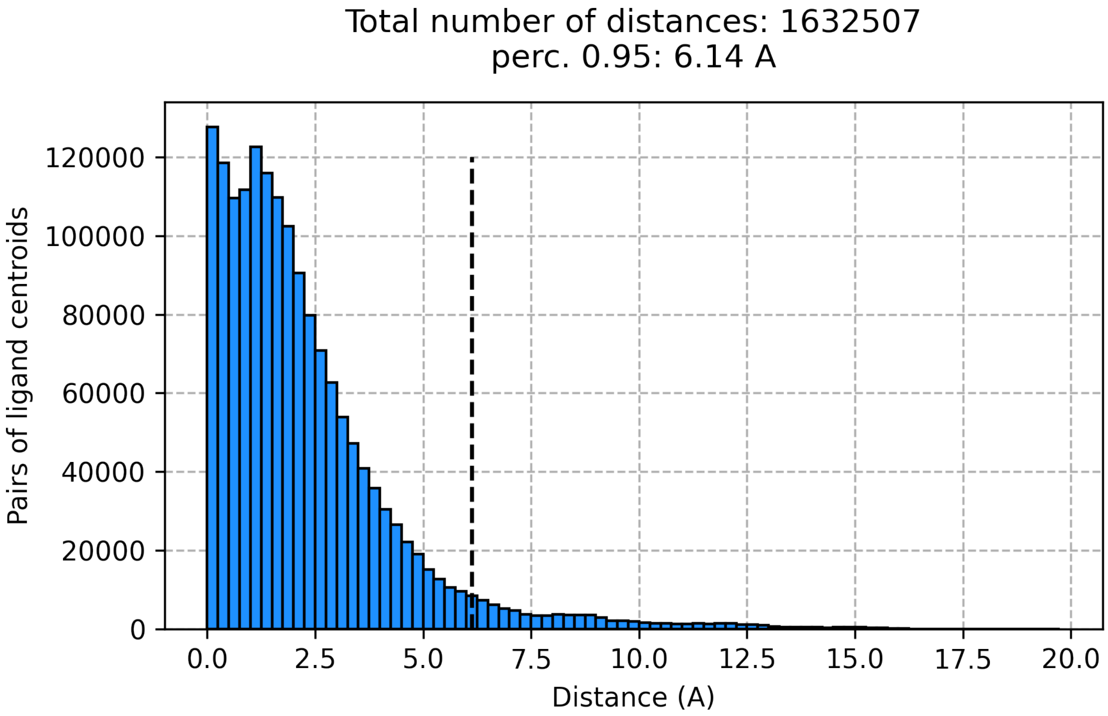


**Fig S25**: Distribution (y-axis) of distances (x-axis) between clustered ligand centroids. In the PDB-LIG set, a total number of 1,604 pockets are identified, encompassing 27,884 ligand centroids. Out of ~1.6M pairwise distances among clustered ligand centroids, only 5% of them are >6.14Å.

| | D1 | D1.2 | D2 | D3 | D4 | D5 | D5.2 | D7 | Av | W. Av. |
|---|---|---|---|---|---|---|---|---|---|---|
| **Number of pairs (k)** | 106.3 | 2 | 108.2 | 26.9 | 26.9 | 6.4 | 10 | 56.4 | - | - |
| **Best Method(s)** | SiteAlign | SiteAlign, KRIPO | SiteAlign | BindSiteS-CNN | SiteAlign | KRIPO | KRIPO | SiteAlign | SiteAlign | SiteAlign |
| **Highest AUROC** | 0.97 | 1.00 | 1.00 | 0.91 | 0.8 | 0.76 | 0.77 | 0.87 | 0.83 | 0.92 |
| **-** | | | | | | | | | | |
| **PocketVec** | **0.97** | **1.00** | 0.96 | 0.67 | 0.72 | 0.64 | 0.62 | **0.87** | 0.81 | 0.89 |
| **Deeplytough** | 0.95 | 0.98 | 0.90 | 0.76 | 0.75 | 0.67 | 0.63 | 0.83 | 0.81 | 0.87 |
| **FuzCav (MOL2)** | 0.94 | 0.99 | 0.99 | 0.69 | 0.58 | 0.55 | 0.54 | 0.77 | 0.76 | 0.86 |
| **FuzCav (PDB)** | 0.94 | 0.99 | 0.98 | 0.69 | 0.58 | 0.56 | 0.54 | 0.77 | 0.76 | 0.86 |
| **SiteAlign** | **0.97** | **1.00** | **1.00** | 0.85 | **0.80** | 0.59 | 0.57 | **0.87** | **0.83** | **0.92** |
| **KRIPO** | 0.91 | **1.00** | 0.96 | 0.60 | 0.61 | **0.76** | **0.77** | 0.85 | 0.81 | 0.86 |
| **TIFP (MOL2)** | 0.66 | 0.90 | 0.91 | 0.66 | 0.66 | 0.71 | 0.63 | 0.71 | 0.73 | 0.75 |
| **TIFP (PDB)** | 0.55 | 0.74 | 0.78 | 0.56 | 0.57 | 0.54 | 0.53 | 0.66 | 0.62 | 0.64 |
| **BindSiteS-CNN** | 0.94 | 0.98 | 0.83 | **0.91** | 0.79 | 0.64 | 0.66 | 0.78 | 0.82 | 0.85 |
| **Median Performance** | 0.94 | 0.99 | 0.94 | 0.69 | 0.64 | 0.62 | 0.60 | 0.78 | 0.78 | 0.85 |
| **Average Performance** | 0.86 | 0.95 | 0.92 | 0.72 | 0.67 | 0.63 | 0.61 | 0.78 | 0.77 | 0.83 |

**Table S1**: Performances (AUROCs) of existing strategies (Deeplytough[5], FuzCav[6], SiteAlign[7], KRIPO[8], TIFP[9], BindSiteS-CNN[10]. File format is specified if needed) to compare pockets based on pocket descriptors/fingerprints among ProSPECCTs datasets. TOP-3 rows include the number of pairs within each dataset together with the best performing method. Median and Average performances have been calculated without considering PocketVec results (shown in blue).

**Table S2**: Binarized kinase-inhibitor matrices from Kleager et al. (2017 set)[3] and Reinecke et al. (2023 set)[4] binarized at 30 nM.
